# Standardized benthic imaging with small, modular ROVs: an open-source framework developed in temperate kelp forests

**DOI:** 10.64898/2026.09.30.755529

**Authors:** Zachary Randell, Megan Williams, Reid Thomson, Keenan Johnson, Clyde McQueen

## Abstract

1. Kelp forests support ecologically, commercially, and culturally important species, yet their benthic communities remain difficult to monitor consistently across space and time. Scientific SCUBA surveys provide valuable observations but can be constrained by safety, bottom time, depth, and the spatial extent that divers can efficiently cover, motivating complementary approaches for collecting standardized benthic imagery.
2. We developed and field-tested a standardized, open-source framework for benthic visual-imaging surveys using a small, modular remotely operated vehicle (ROV). Across four cost-scalable hardware configurations, we integrated navigation sensors, custom BlueOS Extensions, and our surftrak contribution to ArduSub, which enables fixed-altitude flight above the seafloor, and developed two open-source desktop applications that streamline field operations (rov flight ops) and imagery processing (rov imagery processing). We also evaluated power supplies, benthic lighting, camera settings, field protocols, and telemetry-processing workflows over 303 transects conducted on 64 survey days from 2022 to 2026.
3. The ROV operated across soft-sediment and rocky habitats, currents of 0–3 kt, and underwater visibility of 0–10 m. surftrak held the ROV within *±*0.2 m of its target altitude for 96% of 21 h of survey transects, producing an approximately 1 m^2^ standardized image footprint. Sufficient illumination and optimized camera settings yielded high-resolution imagery in visibility below 2 m, while topside power increased endurance and operational flexibility. Combining Doppler velocity log and underwater acoustic positioning data through the vehicle’s extended Kalman filter improved navigation and post hoc transect reconstruction. During 2026 surveys, rov flight ops and rov imagery processing automated flight logging, log retrieval, fault diagnosis, and the sorting, development, and telemetry annotation of survey imagery. Dense kelp canopies nevertheless constrained vehicle operation, and downward-facing imagery provided limited observations of cryptic or obscured organisms.
4. We provide configuration-specific bills of materials, hardware and wiring documentation, software and firmware inventories, camera settings, open-source field and imagery-processing software, step-by-step field protocols, and pre- and post-dive checklists to support replication and adaptation of the framework. Together, these resources provide a field-tested starting point for expanding standardized benthic imaging while complementing diver surveys across kelp forests and other shallow coastal ecosystems.

## Introduction

Kelp forests are highly productive systems that support diverse communities and generate ecological functions and services both locally and through meta-ecosystem connectivity (Dayton, 1985; Steneck *et al*., 2002; Edwards, 2025). Consequently, they are intensively monitored for their importance to fish, invertebrates, seabirds, and marine mammals of commercial and conservation concern, including species listed as threatened or endangered at local or national levels (Eger *et al*., 2023). Canopy-forming kelps that reach the sea surface create floating habitat visible from above and are critical to fish and other taxa during sensitive life stages (Carr, 1991; Johnson, 2006). Likewise, the understory algae and associated benthic communities along the seafloor provide food, shelter, and recruitment habitat for numerous fishes and invertebrates (Duggins *et al*., 1990; Pérez-Matus *et al*., 2025). However, kelp forests are increasingly challenged by warming waters, shifts to alternative states dominated by herbivorous sea urchins or turf algae, the spread of invasive algae, and the absence, extirpation, or disease-induced loss of keystone consumers whose trophic effects support kelp and associated species (Filbee-Dexter & Wernberg, 2018; Rogers-Bennett & Catton, 2019; Galloway *et al*., 2023).

Despite their importance, kelp forests remain difficult to monitor, particularly along the benthos. Surface canopies can often be mapped from air using drones, crewed aircraft, or satellites (Pfister *et al*., 2018; Bell *et al*., 2023), but the vast majority of kelp species and other brown, red, green, and coralline algae, fish, invertebrates, and substrate are only visible underwater along the benthos. Consequently, subtidal kelp forest monitoring has mainly relied on scientific SCUBA diver surveys, often using strip/band transects and point counts (Freiwald *et al*., 2021). These long-term diver-based programs have generated foundational insight into kelp forest dynamics, though surveys are constrained by safe-operation logistics, depth, and the spatial extent that can realistically be covered (Levin, 1992; van Rein *et al*., 2009).

Marine visual imaging complements diver-based surveys and is increasingly used to help establish and monitor marine protected areas, conduct baseline ecological surveys, and support a range of exploratory and applied monitoring efforts (Mallet & Pelletier, 2014). Image-based surveys provide a permanent, revisitable record of fine-scale biological and physical patterns along the benthos, allowing taxa, substrate, and habitat structure to be annotated and quantified consistently, provided imagery is of sufficient quality and resolution (Bowden *et al*., 2020). Downward-facing imagery taken at a consistent camera altitude standardizes the seafloor area captured per photograph, allowing observations to be expressed as densities, percent cover, or other area-normalized metrics and compared among images, transects, sites, and survey periods, while overlapping images can support orthomosaics or three-dimensional reconstructions across larger areas (Pizarro *et al*., 2017).

These approaches have been especially effective in clear-water systems such as coral reefs, where diver and remotely operated vehicle (ROV) methods have provided deep insight through photogrammetry and ecological monitoring of the benthos (Burns *et al*., 2015; Bayley & Mogg, 2020). However, temperate rocky reefs, such as kelp forests, are more difficult to image. Dark water, turbidity, and macro particulates reduce image quality and complicate the collection of standardized benthic imagery, particularly when illumination must be carried by the survey platform itself (Spyksma *et al*., 2022; Bell *et al*., 2026). In these settings, low visibility, ROV-mounted illumination, and variable benthic terrain make consistent ROV altitude essential: flying too high can reduce resolution and illumination, whereas flying too low can result in uneven lighting and motion blur.

While ROVs can expand benthic visual-imaging surveys across greater kelp forest spatial extents and depths than divers can efficiently sample, they have not been systematically integrated into most kelp forest monitoring programs. Historically, many ROV systems were designed for deepsea exploration, offshore inspection, or large-vessel operations, and were therefore too large, too expensive, or too logistically demanding for routine nearshore ecological work from small vessels (Capocci *et al*., 2017; Sward *et al*., 2019; Dalhatu *et al*., 2023). More recently, comparatively low-cost, modular, hand-deployable “micro ROVs” have become commercially available, creating a practical opportunity to expand ROV use in shallow coastal ecosystems (Buscher *et al*., 2020; Overrein *et al*., 2024). Yet adapting such systems for rigorous ecological surveys in kelp forests introduces several challenges. First, as noted above, seasonal upwelling, nutrients, and terrestrial runoff often create dark, turbid water that limits image quality. Second, canopy-forming kelp create vertical structure that can readily entangle a tethered vehicle (Yoklavich *et al*., 2015). Third, swell and current place strong demands on flight control, vehicle power, and pilot workload, especially when surveys require consistent speed, altitude, and image scale (Sward *et al*., 2019).

Additionally, the shallow depths typical of nearshore surveys (*<*30 m, and often 2–15 m) complicate acoustic positioning. In particular, the relatively small angle between the ROV and a vessel-mounted acoustic antenna can reduce the efficacy of Ultra Short Baseline (USBL) positioning systems, especially in rocky habitats where acoustic shadows are common and background positioning noise is already substantial (Vickery, 1998; Sun *et al*., 2024). This matters because consistent positioning underpins not only safe and stable flight operations, but also survey area quantification, spatial analyses, and future autonomous or semi-autonomous ROV workflows (Sørensen *et al*., 2025). More advanced positioning systems exist but often cost $50k or more — more than the rest of a small ROV survey platform — which may be impractical for many non-profit, Tribal, academic, and agency users seeking comparatively low-cost nearshore ROV capacity.

Here we describe and evaluate a practical framework for conducting benthic visual imaging ROV surveys in bull kelp (*Nereocystis luetkeana*) forests and other shallow nearshore ecosystems using small, modular, and comparatively low-cost hardware. Using Blue Robotics’ (BR) BlueROV2 Heavy as our base vehicle, we iterated through four hardware, sensor, and software configurations (*V* 1–*V* 4) that provided increasingly rigorous and repeatable survey capability, and make four contributions: (*1*) surftrak, a terrain-following flight mode that we developed and that has been part of ArduSub since version 4.5 (McQueen, 2025a; ArduPilot Development Team, 2024), which holds the ROV at a fixed altitude above the seafloor to standardize the image footprint; (*2*) hardware modifications and open-source BlueOS Extensions that integrate a Doppler Velocity Log (DVL), USBL acoustic tracking system, and satellite compass for real-time navigation and post hoc spatial reconstruction; (*3*) a cost-scalable evaluation of the power supplies, benthic lighting, camera settings, and field protocols needed for high-resolution, georeferenced photographs in dark, turbid, temperate coastal waters; and (*4*) two open-source desktop applications, rov flight ops and rov imagery processing, that shepherd a survey from pre-dive checks to analysis-ready imagery with less time, training, and manual effort. Although developed in kelp forests, these components address constraints shared by shallow benthic imaging surveys generally — low visibility, variable terrain, and shallow-water positioning — and, because the BlueROV2 is a widely adopted platform with an active open-source community, each can be adopted incrementally, modified for local conditions, and improved through continued development. For kelp forest monitoring, we present this framework not as an alternative to divers, but as a complementary window of observation that can expand benthic survey capacity and provide a practical starting point for ecologists, conservation practitioners, and other coastal users interested in adapting micro ROVs for research, monitoring, and management.

## Methods

Use of AI tools: The rov flight ops and rov imagery processing desktop applications described here were developed, and the manuscript text was edited, with assistance from Anthropic’s Claude (Claude Code; Opus 5.5, 2025–2026). All AI-assisted code was reviewed, covered by automated tests, and field-tested by the authors; the corresponding author takes responsibility for the software and text.

### Overview

Since 2022, the Seattle Aquarium (Aquarium) has iterated through multiple ROV configurations for ecological surveys, with particular emphasis on benthic, downward-facing illumination and camera settings; power supply; vehicle size; and the sensor hardware and software suite needed to improve ROV positioning and support georeferencing of imagery. Between August 2022 and September 2026, we refined this framework across 303 ROV transects on 64 survey days in Elliott Bay, central Puget Sound, the San Juan Islands, and the western Strait of Juan de Fuca (flight log in the project repository). To make this development process useful to other entities seeking a similar cost-scalable micro ROV program, we structure the Methods and Results/Discussion around four hardware iterations, *V* 1–*V* 4 (Table 1; see also Table S4 for core ROV components). Because our primary objective was to obtain high-quality benthic imagery, we organized *V* 1–*V* 4 around two design axes: (*1*) benthic illumination and (*2*) ROV power supply. These two factors most strongly influenced image quality, system cost, and survey endurance, and therefore constrained each version’s broader mission suite. Below, we describe the core *V* 1–*V* 4 ROV hardware, power supply, and lighting and imaging systems; the sensors and topside hardware used for positioning; and the field and imagery software that links them into a single survey workflow.

**Table 1:** Four ROV iterations tested between 2022 and 2026, with ROV power and benthic illumination the two primary determinants of version, and practical recommendations for matching configuration to survey objective. ROV power was supplied either by the standard 14.8 V 18 Ah BR battery or by the Outland Topside Power Supply (OTPS) and a 400 Ah topside battery bank on the survey vessel. Costs are for the vehicle with no navigation sensors (Base), with a DVL (+DVL), and with the DVL, modified USBL, and GNSS compass (Full suite). Add USBL alone for approximate real-time navigation to known survey sites; add DVL alone for fixed-altitude, area-normalized imagery via surftrak; combine DVL, USBL, and ArduSub’s EKF for real-time navigation and post hoc reconstruction of georeferenced transects. Costs exclude the topside laptop and other peripheral equipment; see Tables S3 and S4 for a full *V* 1–*V* 4 hardware breakdown.

| Iteration | ROV power | Benthic lights (lm) | Capability added & recommended use | Approx. cost (k USD) |  |  |
| --- | --- | --- | --- | --- | --- | --- |
|  |  |  |  | Base | +DVL | Full suite |
| V1 | Battery | BR Lumen Lights 6k | Baseline BlueROV2 with downward-facing video and qualitative benthic observation — ideal for scouting, inspection, and rapid deployment. | 9 | 18 | 31 |
| V2 | Battery | Kraken 58k | High-output external lighting and raw photography for sharp benthic imagery — ideal for photo surveys in calm conditions. | 11 | 20 | 33 |
| V3 | OTPS | Kraken 58k | Surface-supplied ROV power for longer field days and dynamic-water operation — ideal for extended photographic transects. | 28 | 37 | 50 |
| V4 | OTPS | DeepSea 40k | Surface-powered lighting integrated with vehicle power — ideal for repeated, extended surveys needing maximum control and endurance. | 37 | 45 | 58 |

### ROV core hardware, power supply, and benthic lighting and imaging

The Aquarium operated two BlueROV2s: ROV *Nereo* (*V* 2), a battery-powered backup vehicle, and ROV *Lutris* (*V* 4), the primary survey vehicle with surface-supplied power and integrated lights. See Table S2 in the Supplementary Material for all ROV software and firmware and Table S4 for all primary ROV hardware. *V* 1 was the smallest configuration and did not include the BR payload skid (Fig. 1a, d), whereas the larger light systems used in *V* 2–*V* 4 utilized the payload skid (Fig. 1b, e, c, f), which added 19.7 cm to the vehicle height, totaling 45.1 cm. Modifications and additions to the ROV mitigated the risk of physical entanglement from kelp stipes and other vertical obstructions in the water column (Section S5).

**Figure 1:**
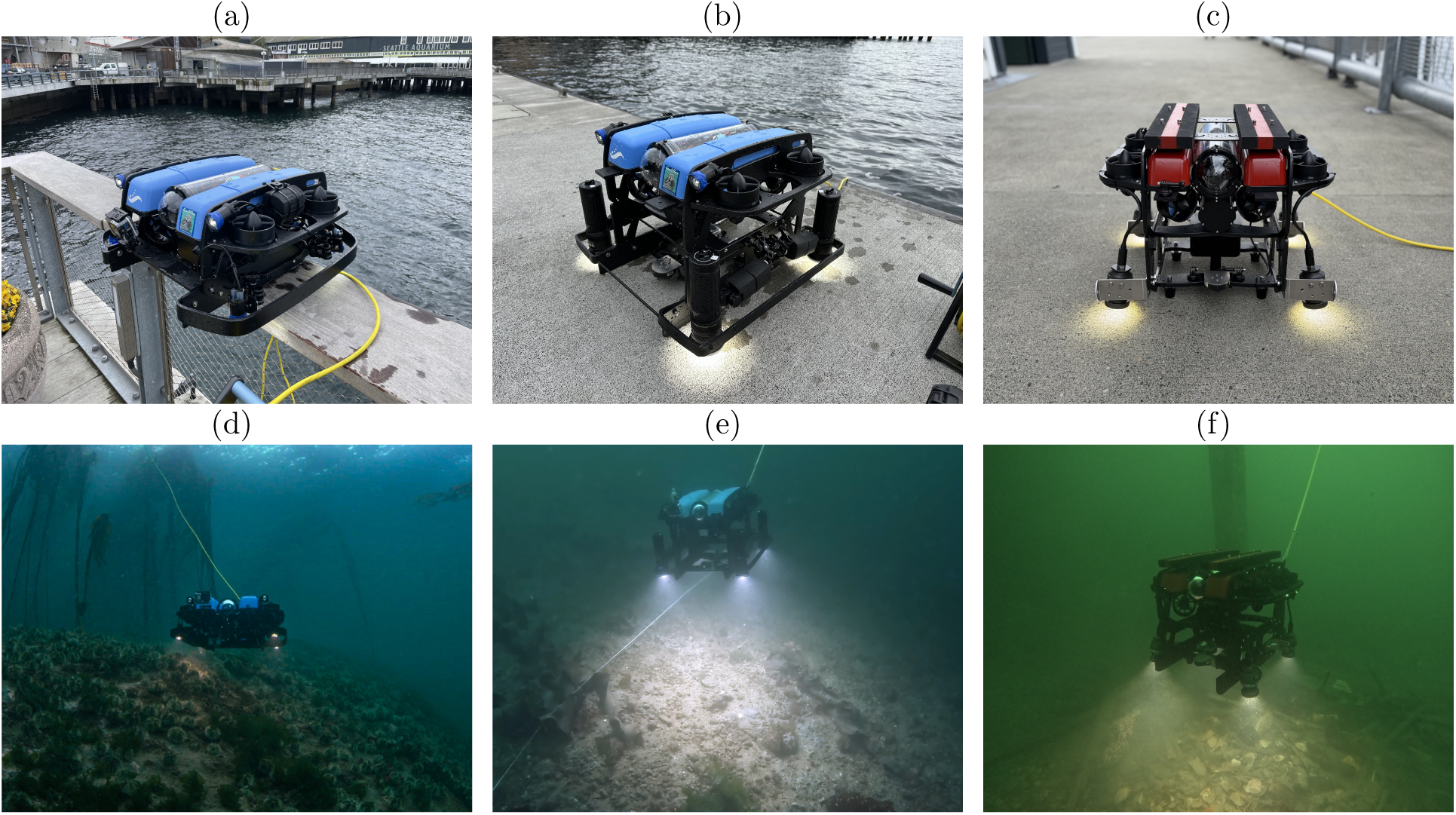
(a), (d) *V* 1: ROV *Nereo* with four integrated BR lights — the simplest, least expensive, and most readily reproducible version. (b), (e) ROV *Nereo* upgraded to a battery-powered *V* 2 configuration, and ROV *Lutris* underwater in *V* 3 with surface-supplied power; both utilize four Kraken lights, with “kelp guard” HDPE entanglement protection visible around the lights in *V* 1– *V* 3. (c), (f) ROV *Lutris* in the *V* 4 configuration with DeepSea lights powered via the OTPS system, across a narrower benthic field of view.

**Figure 2:**
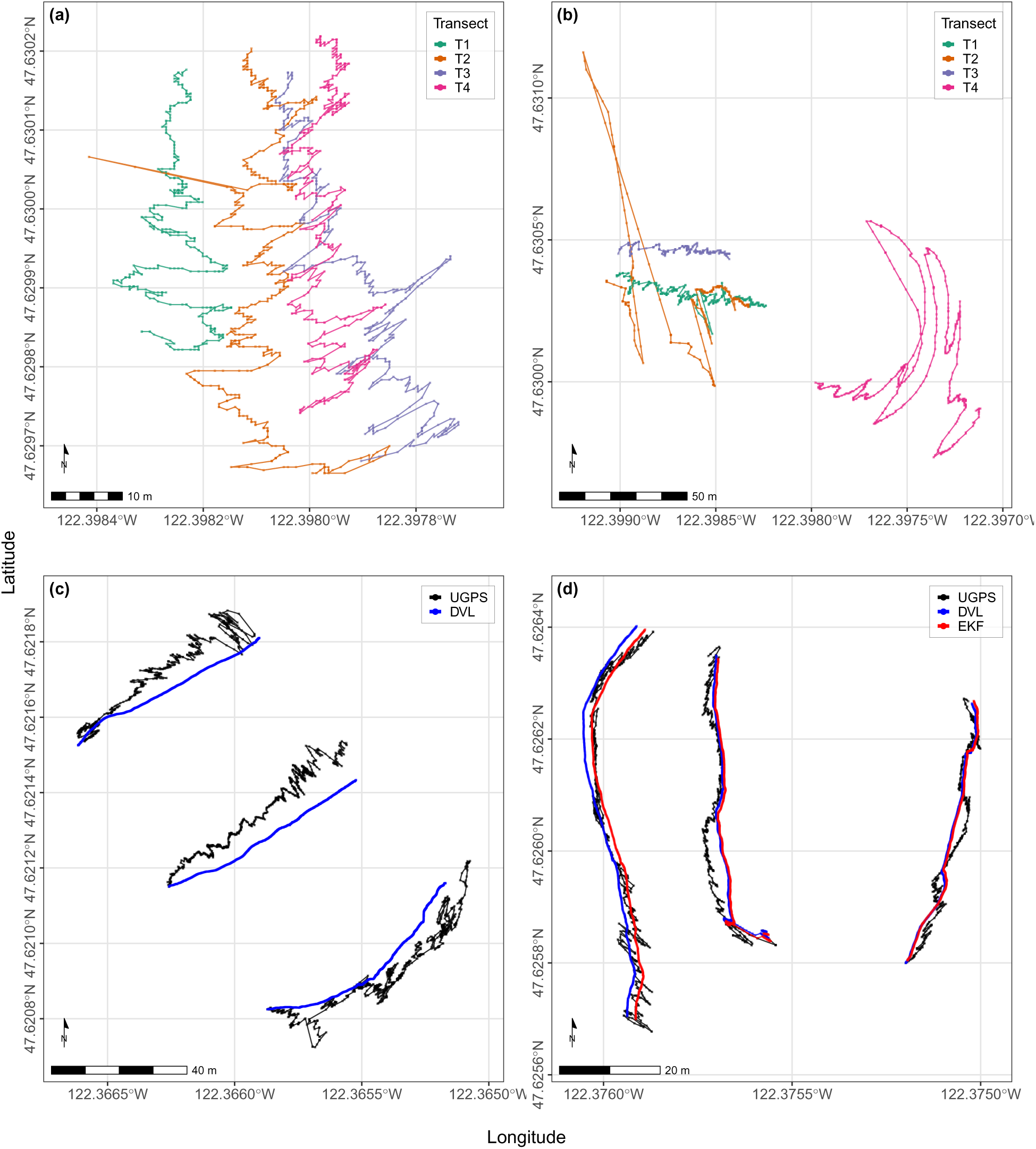
ROV positioning tracking across improvements to hardware and software. (a) Four parallel 100 m transects recorded 2023-03-30, showing consistently noisy UGPS positioning. (b) Four parallel 100 m transects recorded 2022-10-06, showing noisy but generally correct positioning in two transects and two incidents of wild, erratic swings from USBL error. (c) After USBL hardware modifications, a GNSS satellite compass, and BlueOS Extension software, USBL performance improved (black) though not yet fused with DVL data (blue) via ArduSub’s EKF. (d) UGPS (black) and DVL (blue) data fusion via the EKF now generates consistently smooth, accurate tracks (red).

The standard BR 14.8 V 18 Ah battery effectively powered *V* 1 and *V* 2, but multi-hour survey days required two or three batteries. For extended operations with *V* 3 and *V* 4, we therefore installed a topside power system on the Aquarium’s 24 ft aluminum Almar vessel: two 200 Ah lithium-ion batteries and a 2000 W inverter (Fig. S4) feeding the Outland Technology Topside Power Supply (OTPS), which steps voltage up for efficient transmission down the tether and back down at the ROV busbar, delivering a maximum instantaneous output of 1000 W (Fig. S5).

We tested three benthic lighting configurations. To minimize backscatter — the illumination of particles in the water column — all iterations of lights were mounted as far from the downward-facing camera sensor as possible while still minimizing a compact vehicle footprint. For *V* 1, we used four daisy-chained BR lights, each providing 1500 lumens (lm) with a 135*^◦^* beam angle for a total downward-facing output of 6000 lm (Figs. 1a, d). To optimize camera settings for improved image quality, *V* 2 and *V* 3 added four battery-powered Kraken Solar Flare dive lights (Figs. 1b, e). Each light was capable of 18,000 lm with a 125*^◦^* beam angle, though we ran them at 80% for a total output of 57,600 lm and an approximate run time of 70 min. For *V* 4, to increase lighting control, reduce beam angle, and eliminate the need to replace Kraken batteries between flights, we integrated four DeepSea LED SeaLite 2075 SeaStrobes into the ROV using a custom 1 L aluminum BR enclosure mounted on the lower payload skid (Figs. 1c, f). At 100% power output each SeaLite provided 10,000 lm with a 75*^◦^* beam angle. To control the lights, we developed a custom PWM-based control system that enabled pilot-adjustable brightness in 10% increments via the same signaling pathway used by the BR lights. See Section S5 for the control-system design, Table S3 for a bill of materials (BOM), and Figs. S7 and S8 for images and a wire diagram.

Cameras were mounted either directly on the ROV’s high-density polyethylene (HDPE) frame or on the payload skid. For *V* 1, a GoPro HERO 10 was used to collect downward and forward-facing 4K videos. *V* 2–*V* 4 utilized the additional benthic lighting to shoot downward-facing 27.3 MP raw (GPR) photographs using GoPro HERO 12 cameras. Photographs from a transect (typically 50– 200 images) were batch edited together in Adobe Lightroom Classic. See Section S6 for additional details, camera settings, and *V* 1–*V* 4 ROV survey imagery.

### ROV sensors, positioning, and topside configuration

We combined multiple comparatively low-cost sensors, developed BlueOS Extensions to integrate hardware and improve ROV positioning, and enabled semi-autonomous flight control via surftrak (“surface tracking”) terrain hold. The complete navigational suite — including the DVL, USBL, and satellite compass — was compatible across *V* 1–*V* 4. Because consistent downward-facing imagery depends on maintaining a stable altitude above the seafloor, we used both a BR Ping1D and a Water Linked DVL A50 to estimate ROV altitude, with the DVL retained across *V* 1–*V* 4 for the additional precision of its four acoustic beams, whose raw data we log via a modified BlueOS Extension (Section S3). surftrak (McQueen, 2025a; ArduPilot Development Team, 2024) enables the ROV to autonomously maintain a pilot-defined altitude by monitoring range finder messages from either the Ping1D or the DVL.

We also modified Water Linked’s Underwater GPS (UGPS), a USBL acoustic positioning system, to reduce error and improve both real-time navigation and post-survey mapping. We rigidized its acoustic antenna and mount, and replaced the GPS receiver and compass in its topside G2 box with an Advanced Navigation GNSS satellite compass, whose position and heading messages are forwarded to the G2 API by our BlueOS Extension wl ugps external (McQueen, 2025b). See Section S3 for details, Figs. S3–S5 for wiring and topside hardware, and Table S2 for all software and BlueOS Extensions.

### ROV field operations

Most surveys took place in August–October, when annual kelps (Order Laminariales) were still present following the summer algal blooms that degrade water clarity, whereas substrate-focused surveys were conducted in January–March, when annual algae were absent. At minimum, two people were required for flight operations: a pilot and a tether operator, with the tether-operator role trainable in situ by Aquarium volunteers or non-research staff; a Primary Boat Operator was also required, though in calm conditions the same person could serve as both boat operator and pilot. After anchoring at the survey location, we either motored the ROV along the surface to the target depth using the DVL altitude message, or — in higher current — descended directly to the seafloor and navigated in using the USBL positioning system. Once in position, we initialized surftrak, locked the ROV altitude at 0.8 m above the seafloor, and began the transect at a forward speed of 0.10–0.15 m s*^−^*^1^, typically using 20–30% thruster gain in 0–1 kt of current and 40–70% gain in the relatively rare cases of currents up to 3 kt. Transects were conducted parallel to shore (5, 10, 15, 20 m depth) or perpendicular to shore (up a depth gradient, approximately 16–1 m) and ranged 30–100 m in length depending on site conditions and study objectives. Where bull kelp was present, we minimized entanglement risk with simple in-and-out transects: flying into the bed with minimal turning and retracing the same flight path on exit. See Section S2 in the Supplementary Material for complete field protocols, including ROV setup, deployment, and kelp forest entanglement mitigation, and Fig. S1 for the one-page field log used on each survey day. We evaluated performance from 1 Hz transect telemetry: altitude control as the proportion of time within *±*0.10 and *±*0.20 m of the surftrak target altitude logged by the autopilot (Section S3); positioning by comparing USBL, DVL, and fused tracks with one another and with transects flown along 100 m tapes; and power as instantaneous draw and cumulative energy use.

### Field operations and imagery processing software

To reduce the time, expertise, and manual effort required to move from launching the ROV to analysis-ready imagery, we developed two open-source, Python-based desktop applications for the topside laptop (Section S4). rov flight ops is organized as tabs that follow a survey day, with all data filed into a single flight folder (Fig. S6). It (*1*) automatically records topside laptop and tether health, all autopilot parameters, and software versions whenever the vehicle is armed; (*2*) maps the ROV’s live position over an offline chart with survey plans, line-following guidance, and a navigation-sensor readiness check; (*3*) logs transect start and end times, including mid-transect pauses, and previews them against the dive profile; (*4*) downloads, checksum-verifies, and safely clears telemetry logs and imagery from the ROV’s onboard computer; and (*5*) generates a diagnostic flight report and per-transect telemetry CSVs and maps. rov imagery processing then uses the same transect times to (*1*) import GoPro photographs directly from the SD card into per-transect folders, optionally selecting one photograph per user-defined distance traveled (e.g., 1 m); (*2*) batch-develop raw photographs in Adobe Lightroom Classic (crop, chromatic-aberration removal, and AI denoise); and (*3*) trim transect video and composite it with telemetry and forward-camera footage. Both applications supersede our earlier telemetry scripts and read both legacy tlog files and the mcap recordings written by current BlueOS releases.

## Results and Discussion

### ROV flight operations

We successfully conducted ROV surveys from anchored vessels in calm, windy, and rainy conditions, in water currents ranging 0–3 kt, underwater visibility ranging 0–10 m, and across soft-sediment, cobble/pebble, rocky reef, and breakwater riprap habitats. As the ROVs weighed 12.2 kg (*V* 1), 21.1 kg (*V* 2 and *V* 3), and 18.0 kg (*V* 4), one or two people could readily lift, deploy, and retrieve the vehicle without a davit.

The default stabilize and alt hold (“altitude hold”, i.e., depth hold via a barometer sensor) flight modes, combined with the two additional vertical thrusters of the BR Heavy Retrofit Kit, gave the vehicle stable, precise control for benthic transects and enhanced surftrak performance. For benthic surveys, the ROV should be slightly positively buoyant: in alt hold or surftrak, this reduces the need for sustained downward thrust — minimizing thruster wash over soft sediment — while also providing a margin of safety to recover the vehicle in the event of a power loss or other failure. Before surftrak, maintaining a consistent forward speed of 0.10–0.15 m s*^−^*^1^ required continuous monitoring of DVL-derived velocity and near-constant feathering of the thrusters (typically 20–30% gain); once implemented, surftrak shifted the pilot’s primary cognitive load from altitude to forward-velocity management, motivating a proposed ArduSub expansion to autonomously maintain forward velocity as well.

surftrak was highly effective at maintaining a consistent ROV altitude during benthic transects. On a transect flown perpendicular to shore from 16 to 1 m depth (Fig. 3a), a kernel density of realized ROV altitude relative to surftrak’s altitude set point (Fig. 3b) showed tight concordance around the 0.8 m target altitude, with a smaller secondary mode near 0.9 m from when the vehicle had to catch up to steeper or more abrupt changes in topography. This slight lag over steep or complex terrain versus the otherwise smooth tracking across gradual depth change is also visible in the insets of Fig. 3a. Across 118 transects flown in surftrak on 27 survey days in 2024–2025 (20.8 h of 1 Hz telemetry), realized altitude was within *±*0.10 m of the target altitude logged by the autopilot for 85% of the time and within *±*0.20 m for 96%, with a mean absolute error of 0.06 m and no systematic bias (Table S1; Fig. S2). Because image footprint area scales with the square of altitude, *±*0.10 m corresponds to approximately *±*25% in seafloor area per image, which the per-second footprint estimates in the transect CSVs allow to be accounted for when normalizing observations by area. Future development could utilize the DVL’s four logged acoustic beams to anticipate terrain changes before they appear in the averaged altitude signal, or move beyond acoustics entirely toward stereo-camera and computer-vision-based position estimation (e.g., ORB-SLAM3; Campos *et al*., 2021; Marburg *et al*., 2025), which could ultimately support more responsive flight control in complex, high-swell, or rugose environments.

**Figure 3:**
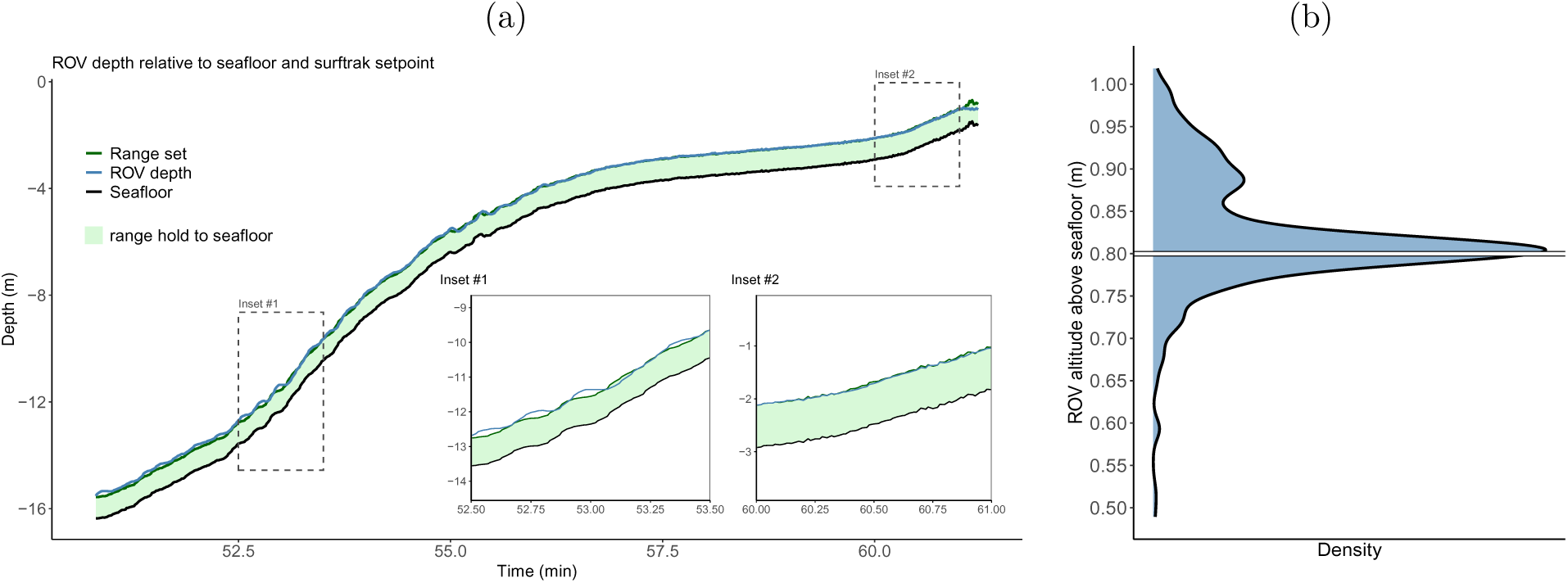
(a) ROV altitude and surftrak performance while flying a transect perpendicular to shore, from 16 to 1 m depth. Tracks show the calculated seafloor (black; ROV depth plus altitude), ROV depth (light blue), and surftrak’s altitude set point (dark green); the light green band marks perfect tracking (seafloor depth plus the set point), so deviation of the ROV depth track from this band indicates straying from the target altitude. Insets show a slight lag catching up to abrupt terrain changes (#1) versus smooth tracking over a gradual gradient (#2). (b) A kernel density of ROV altitude relative to surftrak’s set altitude shows its overall efficacy in maintaining altitude.

Across summers 2022–2025, we completed dozens of transects in low- to moderate-density bull kelp forests in Elliott Bay, Seattle. Bull kelp growing in proximity often becomes tangled such that gaps among stipes (the vertical vegetation that grows up to the sea surface) remain open near the seafloor through which the ROV can pass, but become dense and largely impenetrable higher in the water column; entry was therefore made by flying around bull kelp or through these near-seafloor openings. The highest-risk phase was exiting the bed, as the buoyant tether often rose into the denser tangle of stipes near the surface on the return path; a coordinated tether-tension and lateral-movement technique (Section S2) reliably cleared the tether when this occurred. The ROV was never entangled at depth in a manner that required dive personnel, although there was one instance it became immobilized at the surface in a dense stand of bull kelp blades; we freed the vehicle by pulling anchor, motoring alongside while remaining connected via the tether, and pulling the ROV onto the deck by hand. Our success also reflected conservative risk management: we did not fly into bull kelp denser than we judged passable for the ROV’s height and width. These methods have not yet been tested in giant kelp (*Macrocystis pyrifera*), the dominant canopy-forming kelp in California, and we anticipate highly energetic forests, i.e., with extensive swell, would further challenge ROV operations.

### Power consumption

In the dynamic tidal context of Puget Sound, water current was the primary factor governing both flight operations and power consumption. We therefore targeted slack-high tides whenever possible so that the ROV typically operated in 0–1 kt of current, which proved highly workable. Under these conditions, *V* 1–*V* 3 drew an instantaneous maximum of approximately 100 Watts (W) during benthic transects (see Figs. 4 and S12 for *V* 2–*V* 3 and Fig. S11 for *V* 1), averaging approximately 1 Wh min*^−^*^1^ of cumulative energy use, rounded conservatively. *V* 4’s four integrated DeepSea lights each consumed 100 W at 100% power output, yielding approximately 500 W during standard survey conditions and a cumulative energy consumption of 9.0 Wh min*^−^*^1^ (Fig. S13).

**Figure 4:**
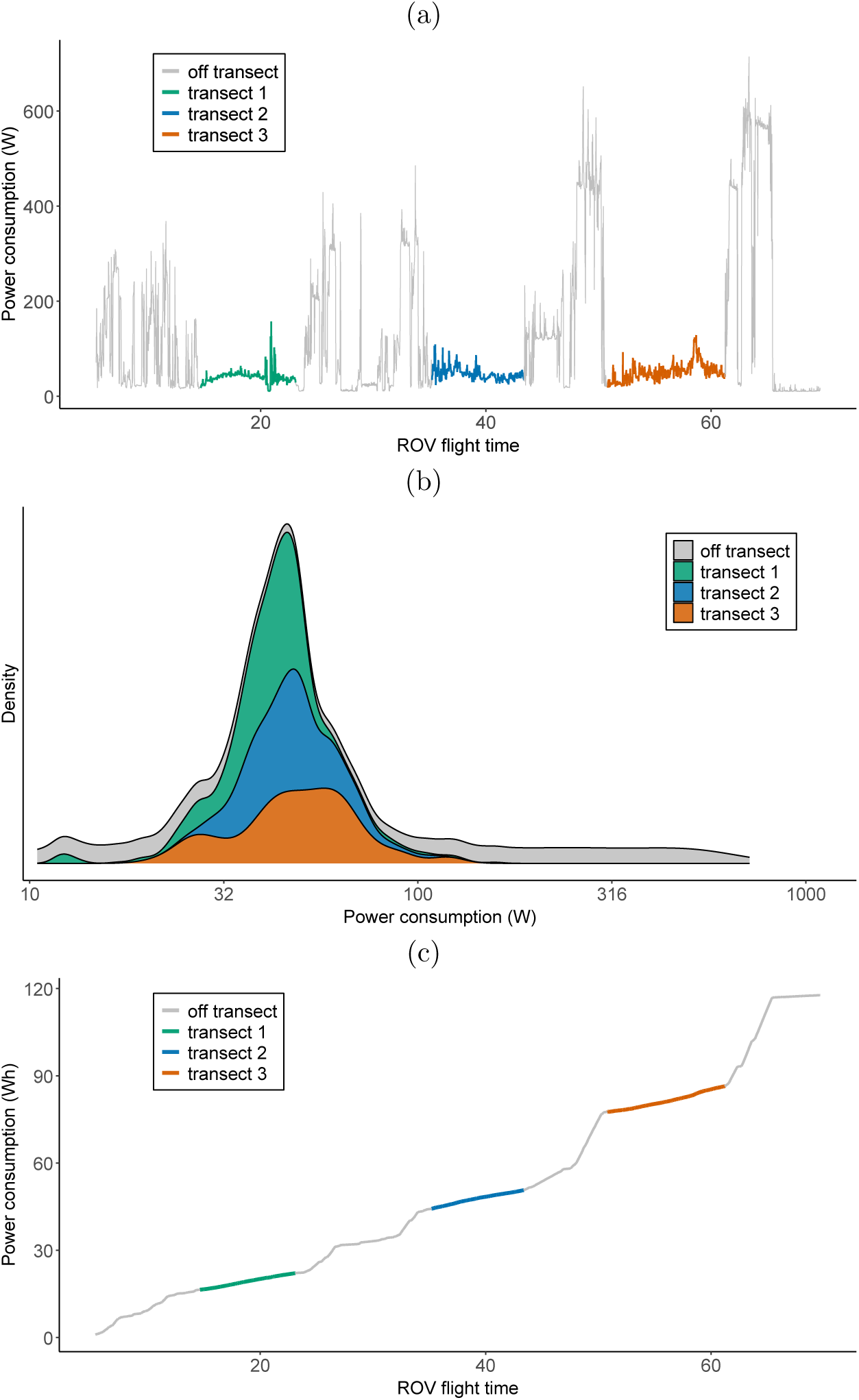
Power consumption for a single contiguous *V* 3 flight in 0–1 kt current in Elliott Bay: (a) instantaneous power (W) over time, with the three transects highlighted in green, blue, and orange and off-transect flight in gray; (b) kernel densities of power draw (log_10_(*x* + 1) transformed; x-axis labels exponentiated back to the natural scale; see Fig. S12 for untransformed data); (c) cumulative power consumption (Wh) for the full 70-minute flight.

Higher currents substantially increased power demand: on a *V* 1 flight in 2–3 kt of current in the western Strait of Juan de Fuca, holding course required sustained thruster output of 50–70%, power rose from approximately 60–100 W to approximately 280 W, and flight time fell to approximately 25 minutes, less than half a typical low-current flight (Fig. S11).

Altogether, the BR battery performed well for *V* 1 and *V* 2 in 0–1 kt current and, where missions do not require back-to-back hours of flight or robust positioning, its simplicity makes it an attractive power solution. Its endurance also matched that of other components: battery-powered GoPro operation was limited to roughly 60 minutes and the Kraken lights (*V* 2, *V* 3) to approximately 70 minutes. However, changing the BR battery required rebooting the ROV, which often also required restarting the topside positioning components (satellite compass, USBL, and associated BlueOS Extensions) — inefficient when survey windows are constrained by fleeting slack tides.

For applications requiring extended flight operations, repeated transects, or extensive work in dynamic water, topside power was a major improvement in endurance and operational flexibility, sustaining elevated thruster output for long periods and eliminating the need to restart the vehicle (and its positioning system) to change batteries. It required substantial up-front investment, however: approximately $12k to professionally install the lithium-ion battery bank and inverter, plus another $13k for the OTPS system itself.

Although *V* 4’s DeepSea lights have yet to be tested in high current, increasing thruster gain from 30 to 50% with the lights at 100% raised consumption by approximately 50 W per 10% step, to approximately 650 W at 50% gain (Fig. S13). Accounting for step-up/step-down conversion losses (approximately 70% efficiency), our 400 Ah topside supply (5120 Wh at 12.8 V) provides an estimated 3584 Wh of usable power — roughly 6.0 hours of runtime at a typical 600 W draw. Altogether, although expensive, topside power may be the single most important investment for users intending to conduct repeated surveys within a day, especially in complex and dynamic environments.

### Positioning

Water Linked’s UGPS system, in isolation, provided an approximate real-time estimate of ROV position relative to the vessel — useful for general situational awareness during flight — but its default configuration was too noisy for rigorous post hoc mapping, with apparent swings of 5–20 m in ROV position that rendered spatial plotting and survey-area quantification largely untenable (Fig. 2a,b). Through repeated testing, we traced this error to movement of the foldable antenna and its mount as the anchored vessel bobbed, and to the GPS receiver and compass in the topside G2 box; rigidizing the antenna and substituting the GNSS compass substantially improved UGPS performance (Fig. 2c; Section S3).

Even so, USBL positioning in shallow water (0–20 m) remained inherently difficult. As the ROV entered very shallow depths (0–10 m), the small angle between the vehicle and the vessel-mounted antenna increased positioning error, and shallow rocky reefs are replete with acoustic shadows that can block the ROV’s signal. USBL’s major strength in these conditions is a real-time, world-referenced estimate of approximate vehicle position; for rigorous benthic mapping or post hoc quantification of survey tracks and area, USBL alone is not sufficient.

By contrast, the Water Linked DVL A50 proved highly valuable for benthic survey operations. Its four downward-facing acoustic beams provided bottom-relative velocity and altitude at approximately 10 Hz, supporting surftrak and precisely recording fine-scale surge, sway, and heave relative to the seafloor. Apart from its cost of approximately $9k, the DVL had two main limitations, both inherent to bottom-referenced navigation without an external world frame: (*1*) limited ability to resolve yaw, such that on out-and-back transects flown along a diver-laid meter tape, small heading errors at the turnaround could rotate the dead-reckoned return path away from the true line even when total distance traveled was captured accurately; and (*2*) no absolute geographic reference, such that even from a known starting position the DVL-estimated path drifted from the true vehicle position over time (Fig. 2c, where blue DVL tracks are offset from the noisier but spatially accurate black USBL tracks). Nevertheless, given the similar prices of the UGPS and DVL and the DVL’s role in enabling surftrak, the DVL would be the clear single-sensor purchase.

The greatest positioning performance emerged when USBL and DVL data were fused through ArduSub’s EKF, together with compass and inertial data from the Navigator Flight Controller: the DVL anchored fine-scale movement while the UGPS provided a world-referenced frame that reduced drift and helped correct the DVL’s weaker handling of yaw, consistently yielding improved real-time navigation and usable post hoc transect reconstruction (Fig. 2d). Across 28 passes flown along 100 m tapes, distance integrated from the EKF’s velocity estimate recovered a median 99.3% of tape length, versus 96.3% from DVL odometry alone (33 passes; Section S4). That said, for users without a marine-technology background, the full system is neither simple nor lightweight to operate: it requires Ethernet switches in both the ROV enclosure and command console and a Power-over-Ethernet supply for the GNSS compass (Figs. S3, S5), setup at the start of a field day is time-intensive, and the system occasionally required power cycling to restore message flow across devices. Failures could also be silent: a sensor could be configured, transmitting, and valid yet not used by the EKF. These challenges motivated the monitoring and diagnostic tools built into rov flight ops (see below).

### Benthic lighting and imaging

*V* 1, which combined the BR battery with four integrated BR lights (6000 lm) and required no payload skid, was the smallest, least expensive, simplest, and most rapidly deployable ROV configuration. As such, it was well suited to scouting, qualitative surveys, inspection, aquaculture applications, and other missions where high-resolution photographs and extensive image post-processing were not required — such as identifying conspicuous species (crabs, sea stars, sea urchins, demersal fish) or marine debris. Figs. S9 and S10 provide examples of still images extracted from 4K video. However, we sought to differentiate among functional groups of macrophytes, including red, brown, and green algae, as well as encrusting and articulated coralline algae, and found that sharper imagery was required. Once extracted from downward-facing 4K video, still frames were too degraded to support downstream workflows that automate ecological data extraction (see Section S6 and Fig. S9). After switching to GoPro HERO 12 photographs and custom GoPro Labs camera settings, the four BR lights also proved insufficient to optimize camera ISO and shutter-speed settings for the sharper imagery desired. This prompted the transition to the battery-powered Kraken lights used in *V* 2 and *V* 3, which increased benthic illumination substantially from 6,000 to 57,600 lm. Shooting raw (GoPro GPR) alongside JPEG previews preserved flexibility for post-processing in Adobe Lightroom Classic, which rov imagery processing now automates (Section S6). Although these four Kraken lights were limited to a run time of 70 minutes and each added 1.3 kg and some degree of drag to the vehicle, the improvement in image quality justified these tradeoffs.

With the *V* 2 and *V* 3 Kraken lights, we consistently gathered high-resolution and visually sharp benthic imagery, even in the dark and often turbid waters of coastal Washington (Fig. 5). The resulting photographs resolved fine-scale textures and morphological features among, e.g., bushy, filamentous, leafy, and branched red algae, as well as distinctions between kelps such as sugar kelp (*Saccharina latissima*) and sieve kelp (*Agarum clathratum*), and among substrates such as sand, silt, shell debris, cobble, pebble, and boulder. At a survey altitude of 0.8 m, cropped downward-facing photographs captured an approximately 1.1 *×* 0.95 m field of view, yielding a consistent footprint of roughly 1 m^2^ of seafloor per image. Ecological data extraction from these photographs, its validation (including paired comparison with diver surveys), and statistical analysis are downstream workflows addressed elsewhere; our focus here is the field acquisition framework needed to generate standardized, analysis-ready imagery.

**Figure 5:**
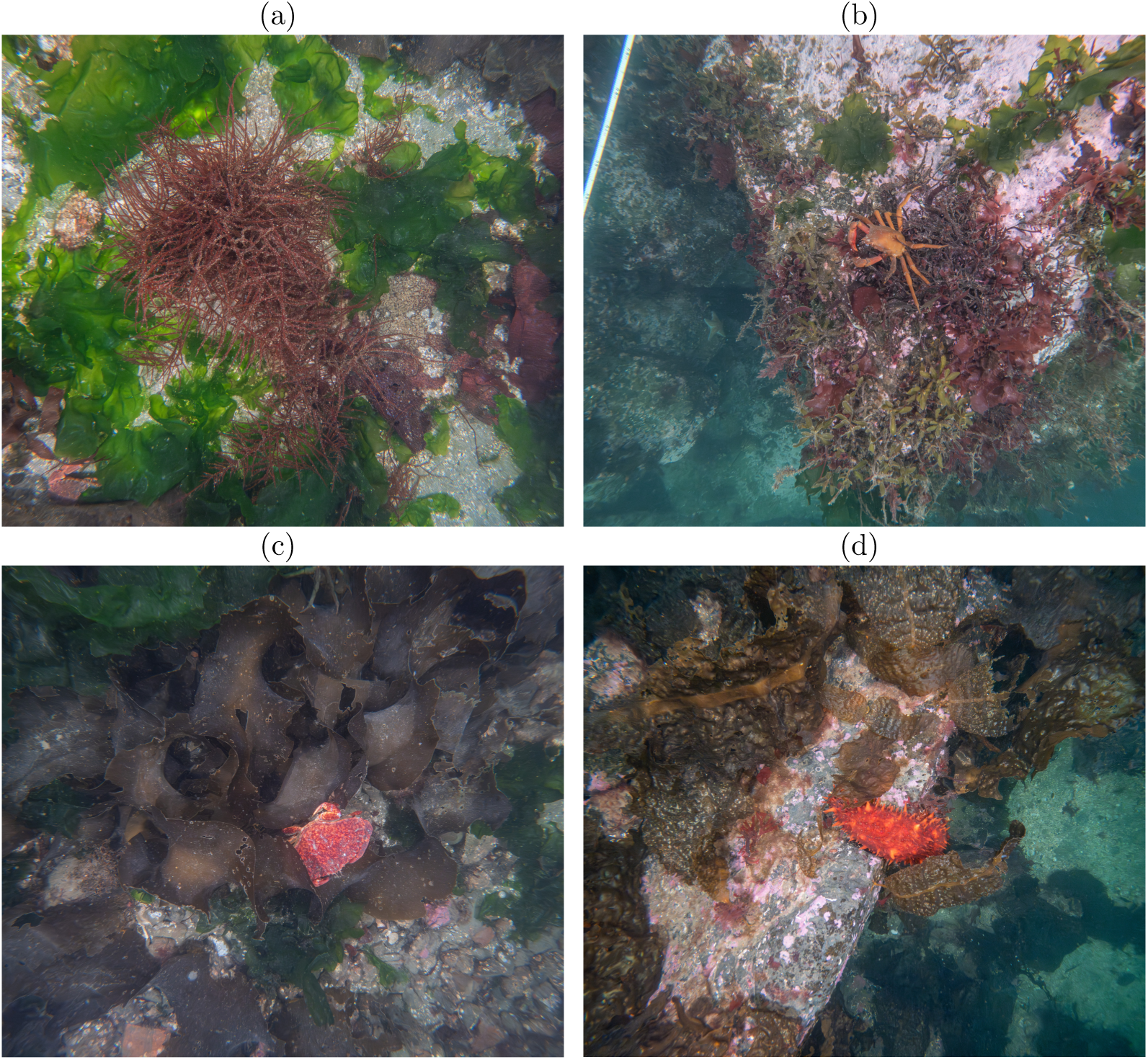
High-resolution, 27.3 MP photos from the *V* 2/*V* 3 Kraken-light configuration, obtained by shooting raw (GPR) and processing in Adobe Lightroom; this level of sharpness is essential for machine-learning-based data extraction. Even illumination across the low-rugosity substrate in (a) and (c) contrasts with (b) and (d), where extensive and moderate seafloor rugosity leaves large or small portions of the image unevenly lit, respectively. See Section S6 for camera settings and additional imagery, and Table S5 for complete transects.

Despite these *V* 2 and *V* 3 improvements in image quality, new lighting challenges emerged. Because each Kraken light had a wide beam angle (125*^◦^*), a substantial fraction of the emitted light fell outside the downward-facing camera field of view. This produced generally even illumination over flat benthos, but sloping seafloor and rugose rocky reef could drop portions of the benthos outside the main illuminated field, reducing local image quality and taxonomic resolution. The fully integrated *V* 4 DeepSea lights addressed this limitation by using a narrower beam angle (75*^◦^*), which concentrated more usable light within the camera field of view and improved illumination efficiency at the target survey altitude. Because these lights are powered and controlled through the ROV, *V* 4 also eliminates separate light batteries and provides a more expandable platform for future camera upgrades, including DSLR-style systems capable of synchronized strobe triggering. Together, these results suggest that qualitative video surveys and quantitative photographic surveys should be treated as distinct mission classes: the former can be supported by simpler configurations (*V* 1), whereas the latter require sufficient illumination, fixed-altitude flight, and camera settings optimized for sharp, standardized benthic imagery (*V* 2–*V* 4).

### Field operations and imagery processing software

We used rov flight ops and rov imagery processing (or, before 13 September, their single-program predecessor) on all six Elliott Bay survey days (nine sites) between 31 August and 18 September 2026, sorting 7,039 raw photographs into 36 transects. Across the six flights recorded by rov flight ops from 15–18 September, the applications logged 6.4 hours of topside telemetry at 1 Hz and downloaded and checksum-verified 31 GB of vehicle logs and imagery. Because transect times entered once now drive every downstream step, tasks that previously required several scripts, manual file handling, and experienced staff can be completed by any trained crew member — all by the end of the survey day.

The applications also exposed failure modes that are otherwise silent. Within minutes of recovery on 17 September, the automated flight report attributed five ground-station failsafe disarms to brief losses of the tether link, four of which preceded the disarm by 1–2 s, and showed that the link was less reliable under Cockpit than under QGroundControl that day — an analysis that had previously taken an evening of manual log review. rov flight ops’s readiness check separates whether each positioning sensor is configured, receiving, valid, and fused into the EKF, a distinction that proved critical after DVL-only dives on 17–18 September navigated normally but logged no geographic coordinates because the EKF origin had not been applied. For imagery, rov imagery processing detected a 34.8 s offset between GoPro and vehicle clocks on a 14 September test flight by cross-correlating the rotation of the downward-facing image with the autopilot’s yaw rate, and its distance-based photo selection is calibrated against the tape transects described above.

## Conclusions

Small, modular ROVs can provide a repeatable means of collecting standardized benthic imagery in kelp forests and other shallow nearshore ecosystems. Using the BlueROV2 as a base platform, we found that survey capability depended on the integration of illumination, power supply, altitude control, positioning, field and imagery software, and field protocols. Simple configurations supported rapid deployment, scouting, and qualitative video surveys, whereas more developed configurations supported fixed-altitude, high-resolution photography and post hoc spatial reconstruction. The *V* 1–*V* 4 framework therefore provides a cost-scalable structure for matching system complexity to survey objectives (Table 1).

Several practical lessons emerged. Sufficient, well-positioned illumination enabled high-resolution imaging in visibility below 2 m, while surftrak held the ROV within *±*0.2 m of its target altitude 96% of the time across 118 transects, standardizing the photographic footprint for area-normalized metrics. Topside power increased endurance and flexibility for repeated surveys, high-output lighting, and dynamic conditions. Shallow-water positioning remained challenging: USBL supported real-time situational awareness, whereas more rigorous transect reconstruction required DVL-supported motion estimates and sensor fusion. The resulting positioning system was effective but operationally complex, which motivated the monitoring, fault-detection, and diagnostic tools built into rov flight ops. Together, rov flight ops and rov imagery processing consolidate flight logging, log retrieval, fault diagnosis, and imagery processing into two open-source applications, reducing the expertise and time needed to go from launch to analysis-ready imagery.

For kelp forest monitoring, this framework complements rather than replaces scientific divers. ROVs can expand visual access across depths and repeatable survey areas, whereas dense kelp canopies, entanglement risk, and cryptic or obscured organisms remain important limitations. Beyond kelp forests, the same *V* 1–*V* 4 framework is well suited to other shallow coastal applications with similar imaging requirements, including soft-sediment and coral reef benthic surveys, photogrammetry mapping missions, and marine debris identification. To facilitate adoption and replication, we provide configuration-specific bills of materials, hardware and software documentation, open-source field and imagery-processing software, camera settings, operational checklists, and step-by-step field protocols.

## Acknowledgments

We appreciate Dr. Shawn Larson and Dr. Erin Meyer for supporting the initiation of this work at the Seattle Aquarium alongside Kathleen Hurley at the Port of Seattle Maritime Environment and Sustainability. We thank the Aquarium’s Dive Program for facilitating successful field operations, particularly Joel Hollander and Alex Tanz. We are grateful to Dr. Aaron Marburg at the University of Washington’s Applied Physics Lab for his collaboration and assistance debugging the integrated *V* 4 lights. We thank Diane Tomecek at Sea Otter Foundation and Trust and Tami Pokorny at the North Pacific Coast Marine Resources Committee for their early support of this work.

## Funding information

This project has been funded wholly or in part by the United States Environmental Protection Agency under assistance agreement PC-01J89501 through the Washington Department of Fish and Wildlife. The contents of this document do not necessarily reflect the views and policies of the Environmental Protection Agency or the Washington Department of Fish and Wildlife, nor does mention of trade names or commercial products constitute endorsement or recommendation for use. This project was also supported by a Service Agreement from the Port of Seattle P-00322850 Urban Kelp Project.

## Conflict of Interest statement

The authors declare no conflicts of interest.

## Data availability statement

All data and code supporting this manuscript, including the rov flight ops and rov imagery processing desktop applications, are available at https://github.com/Seattle-Aquarium/CCR ROV survey methods. Complete transects of survey imagery are available at https://osf.io/9dkp8/. The repository will be archived on Zenodo with a DOI upon acceptance.

## Author Contributions

ZR supervised the research, implemented ROV hardware, developed ROV field methods, analyzed telemetry logs, and wrote the manuscript. MW implemented ROV hardware, developed ROV field methods, developed ROV camera settings, created code and analyzed telemetry logs, processed imagery, and wrote the manuscript. RT implemented ROV hardware, participated in ROV fieldwork, created code and analyzed telemetry logs, processed imagery, and wrote the manuscript. KJ wrote custom BlueOS Extensions, developed the *V* 4 custom lighting solution, participated in fieldwork, and wrote the manuscript. CM wrote custom BlueOS Extensions and the surftrak ArduSub contribution, participated in fieldwork, analyzed telemetry logs, and wrote the manuscript. ZR, MW, and RT used Anthropic’s Claude (Opus 5.5) to iteratively develop and test the rov flight ops and rov imagery processing desktop applications. All authors contributed critically to the drafts and gave final approval for publication.

## Statement on Inclusion

Our study is part of our broader research program that seeks to collaborate extensively with state, Tribal, federal, non-profit, and academic entities. We routinely participate in local conservation, research, and management gatherings to communicate and coordinate our work such that it can contribute as much as possible to advance Washington kelp forest monitoring, conservation, management, and restoration objectives.

## Data and Code Availability

Code and data supporting this manuscript, including both field applications, are available at https://github.com/Seattle-Aquarium/CCR_ROV_survey_methods; complete transects of survey imagery are available at https://osf.io/9dkp8/. The repository will be archived on Zenodo with a DOI upon acceptance.

## Notes

https://github.com/Seattle-Aquarium/CCR_ROV_survey_methods

### Competing Interest Statement

The authors have declared no competing interest.

## References

ArduPilot Development Team (2024). ArduSub release notes, Sub-4.5.0: SURFTRAK flight mode (altitude hold with rangefinder). https://github.com/ArduPilot/ardupilot/blob/master/ArduSub/ReleaseNotes.txt. Accessed 28 September 2026.

Bayley, D. T. I. & Mogg, A. O. M. (2020). A protocol for the large-scale analysis of reefs using structure from motion photogrammetry. Methods in Ecology and Evolution, 11, 1410–1420.

Bell, J. J., Collina, M., Micaroni, V., Strano, F. & Pierotti, M. (2026). Photogrammetry in rocky temperate marine ecosystems: applications, problems and potential. Ocean Ecosystems, 2, 2.

Bell, T. W., Cavanaugh, K. C., Saccomanno, V. R., Cavanaugh, K. C., Houskeeper, H. F., Eddy, N., Schuetzenmeister, F., Rindlaub, N. & Gleason, M. (2023). Kelpwatch: A new visualization and analysis tool to explore kelp canopy dynamics reveals variable response to and recovery from marine heatwaves. PLOS ONE, 18, e0271477.

Bowden, D., Rowden, A., Chin, C., Hempel, S., Wood, B., Hart, A. & Clark, M. (2020). Best practice in seabed image analysis for determining taxa, habitat, or substrata distributions. Tech. Rep. New Zealand Aquatic Environment and Biodiversity Report No. 239, Fisheries New Zealand.

Burns, J., Delparte, D., Gates, R. & Takabayashi, M. (2015). Integrating structure-from-motion photogrammetry with geospatial software as a novel technique for quantifying 3D ecological characteristics of coral reefs. PeerJ, 3, e1077.

Buscher, E., Mathews, D. L., Bryce, C., Bryce, K., Joseph, D. & Ban, N. C. (2020). Applying a low cost, mini remotely operated vehicle (ROV) to assess an ecological baseline of an Indigenous seascape in Canada. Frontiers in Marine Science, 7, 669.

Campos, C., Elvira, R., Gómez Rodríguez, J. J., Montiel, J. M. M. & Tardós, J. D. (2021). ORBSLAM3: An accurate open-source library for visual, visual-inertial, and multimap SLAM. IEEE Transactions on Robotics, 37, 1874–1890.

Capocci, R., Dooly, G., Omerdić, E., Coleman, J., Newe, T. & Toal, D. (2017). Inspection-class remotely operated vehicles—A review. Journal of Marine Science and Engineering, 5, 13.

Carr, M. H. (1991). Habitat selection and recruitment of an assemblage of temperate-zone reef fishes. Journal of Experimental Marine Biology and Ecology, 146, 113–137.

Dalhatu, A. A., Sa’ad, A. M., Cabral de Azevedo, R. & De Tomi, G. (2023). Remotely operated vehicle taxonomy and emerging methods of inspection, maintenance, and repair operations: An overview and outlook. Journal of Offshore Mechanics and Arctic Engineering, 145, 020801.

Dayton, P. K. (1985). Ecology of kelp communities. Annual Review of Ecology and Systematics, 16, 215–245.

Duggins, D. O., Eckman, J. E. & Sewell, A. T. (1990). Ecology of understory kelp environments. II. Effects of kelps on recruitment of benthic invertebrates. Journal of Experimental Marine Biology and Ecology, 143, 27–45.

Edwards, M. S. (2025). Kelp forests under threat and the loss of ecosystem services they provide. Annual Review of Ecology, Evolution, and Systematics, 56, 365–392.

Eger, A. M., Marzinelli, E. M., Beas-Luna, R., Blain, C. O., Blamey, L. K., Byrnes, J. E. K., Carnell, P. E., Choi, C. G., Hessing-Lewis, M., Kim, K. Y., Kumagai, N. H., Lorda, J., Moore, P., Nakamura, Y., Pérez-Matus, A., Pontier, O., Smale, D., Steinberg, P. D. & Vergés, A. (2023). The value of ecosystem services in global marine kelp forests. Nature Communications, 14, 1894.

Filbee-Dexter, K. & Wernberg, T. (2018). Rise of turfs: A new battlefront for globally declining kelp forests. BioScience, 68, 64–76.

Freiwald, J., McMillan, S. M. & Abbott, D. (2021). Reef Check California instruction manual: A guide to monitoring California’s kelp forests, 10th edition. Tech. rep., Reef Check Foundation, Marina del Rey, CA, USA.

Galloway, A. W. E., Gravem, S. A., Kobelt, J. N., Heady, W. N., Okamoto, D. K., Sivitilli, D. M., Saccomanno, V. R., Hodin, J. & Whippo, R. (2023). Sunflower sea star predation on urchins can facilitate kelp forest recovery. Proceedings of the Royal Society B: Biological Sciences, 290, 20221897.

Johnson, D. W. (2006). Predation, habitat complexity, and variation in density-dependent mortality of temperate reef fishes. Ecology, 87, 1179–1188.

Levin, S. A. (1992). The problem of pattern and scale in ecology. Ecology, 73, 1943–1967.

Mallet, D. & Pelletier, D. (2014). Underwater video techniques for observing coastal marine biodiversity: A review of sixty years of publications (1952–2012). Fisheries Research, 154, 44–62.

Marburg, A., Guimond, N., Cheruvathur, G. M., Liao, C., Shao, M., Lalli, T., Heller, P., Randell, Z. & McQueen, C. (2025). Development of a low-cost payload for littoral ROV navigation. In: OCEANS 2025 - Great Lakes.

McQueen, C. (2025a). ardusub surftrak: Add range hold flight mode to ardusub. https://github.com/clydemcqueen/ardusub_surftrak. GitHub repository.

McQueen, C. (2025b). Wl ugps external extension. https://github.com/clydemcqueen/wl_ugps_external_extension. GitHub repository, a BlueOS extension that listens for NMEA messages and sends position information to the Water Linked UGPS system.

Overrein, M. M., Tinn, P., Aldridge, D., Johnsen, G. & Fragoso, G. M. (2024). Biomass estimations of cultivated kelp using underwater RGB images from a mini-ROV and computer vision approaches. Frontiers in Marine Science, 11, 1324075.

Pfister, C. A., Berry, H. D. & Mumford, T. (2018). The dynamics of kelp forests in the northeast Pacific Ocean and the relationship with environmental drivers. Journal of Ecology, 106, 1520– 1533.

Pizarro, O., Friedman, A., Bryson, M., Williams, S. B. & Madin, J. (2017). A simple, fast, and repeatable survey method for underwater visual 3D benthic mapping and monitoring. Ecology and Evolution, 7, 1770–1782.

Pérez-Matus, A., Micheli, F., Konar, B., Shears, N., Low, N. H. N., Okamoto, D. K., Wernberg, T., Krumhansl, K. A., Ling, S. D., Kingsford, M., Navarrete-Fernandez, T., Ruz, C. S. & Byrnes, J. E. K. (2025). Kelp forests as nursery and foundational habitat for reef fishes. Ecology, 106, e70007.

Rogers-Bennett, L. & Catton, C. A. (2019). Marine heat wave and multiple stressors tip bull kelp forest to sea urchin barrens. Scientific Reports, 9, 15050.

Spyksma, A. J. P., Miller, K. I. & Shears, N. T. (2022). Diver-generated photomosaics as a tool for monitoring temperate rocky reef ecosystems. Frontiers in Marine Science, 9, 953191.

Steneck, R. S., Graham, M. H., Bourque, B. J., Corbett, D., Erlandson, J. M., Estes, J. A. & Tegner, M. J. (2002). Kelp forest ecosystems: biodiversity, stability, resilience and future. Environmental Conservation, 29, 436–459.

Sun, L., Wang, Y., Hui, X., Ma, X., Bai, X. & Tan, M. (2024). Underwater robots and key technologies for operation control. Cyborg and Bionic Systems, 5, Article 0089.

Sward, D., Monk, J. & Barrett, N. S. (2019). A systematic review of remotely operated vehicle surveys for visually assessing fish assemblages. Frontiers in Marine Science, 6, 134.

Sørensen, F. F., Mai, C., von Benzon, M., Liniger, J. & Pedersen, S. (2025). The localization problem for underwater vehicles: An overview of operational solutions. Ocean Engineering, 330, 121173.

van Rein, H., Brown, C. J., Quinn, R. & Breen, J. (2009). A review of sublittoral monitoring methods in temperate waters: A focus on scale. Underwater Technology, 28, 99–113.

Vickery, K. (1998). Acoustic positioning systems. A practical overview of current systems. In: Proceedings of the 1998 Workshop on Autonomous Underwater Vehicles.

Yoklavich, M., Reynolds, J. & Rosen, D. (2015). A comparative assessment of underwater visual survey tools: Results of a workshop and user questionnaire. Tech. Rep. NOAA Technical Memorandum NMFS-SWFSC-547, National Oceanic and Atmospheric Administration.

