## Supplementary Material for "Standardized benthic imaging with small, modular ROVs: an open-source framework developed in temperate kelp forests"

### *Supplementary Material:* A modular, open-source framework for benthic ROV surveys in temperate kelp forests

**S6 ROV camera settings and survey imagery** **31**

#### Field log and checklist

We print one field log (Fig. S1) for each survey day and fill it in by hand on deck. It records the day's metadata and the ArduSub, BlueOS, and Cockpit versions; opens and closes each flight with pre- and post-flight checklists that condense the protocols in Section S2 and the `rov_flight_ops` workflow (Section S4); and holds, for up to six transects, the TC 25 start and end times and the target depth, altitude, speed, light level, and flight mode. The transect times are then entered into `rov_flight_ops`'s Transects tab (Section S2), which uses them to subset the telemetry and imagery. The editable source (`field_checklist/`) is in the repository.

Every ROV survey flight from August 2022 to September 2026 is listed in the repository's flight log (`survey_log/`): date, program, site, transect, vehicle, and transect start and end times, with each row marked as counted or not as a survey transect and why (e.g., test and outreach flights, aborted transects). A script in the same folder reproduces the totals given in the main text: 303 transects on 64 survey days, where a survey day is a calendar date on which at least one survey transect was flown. On diver–ROV comparison days, the ROV's passes along each 100 m tape were processed as three 30 m transects to match the divers' Reef Check transects, and are counted as such.

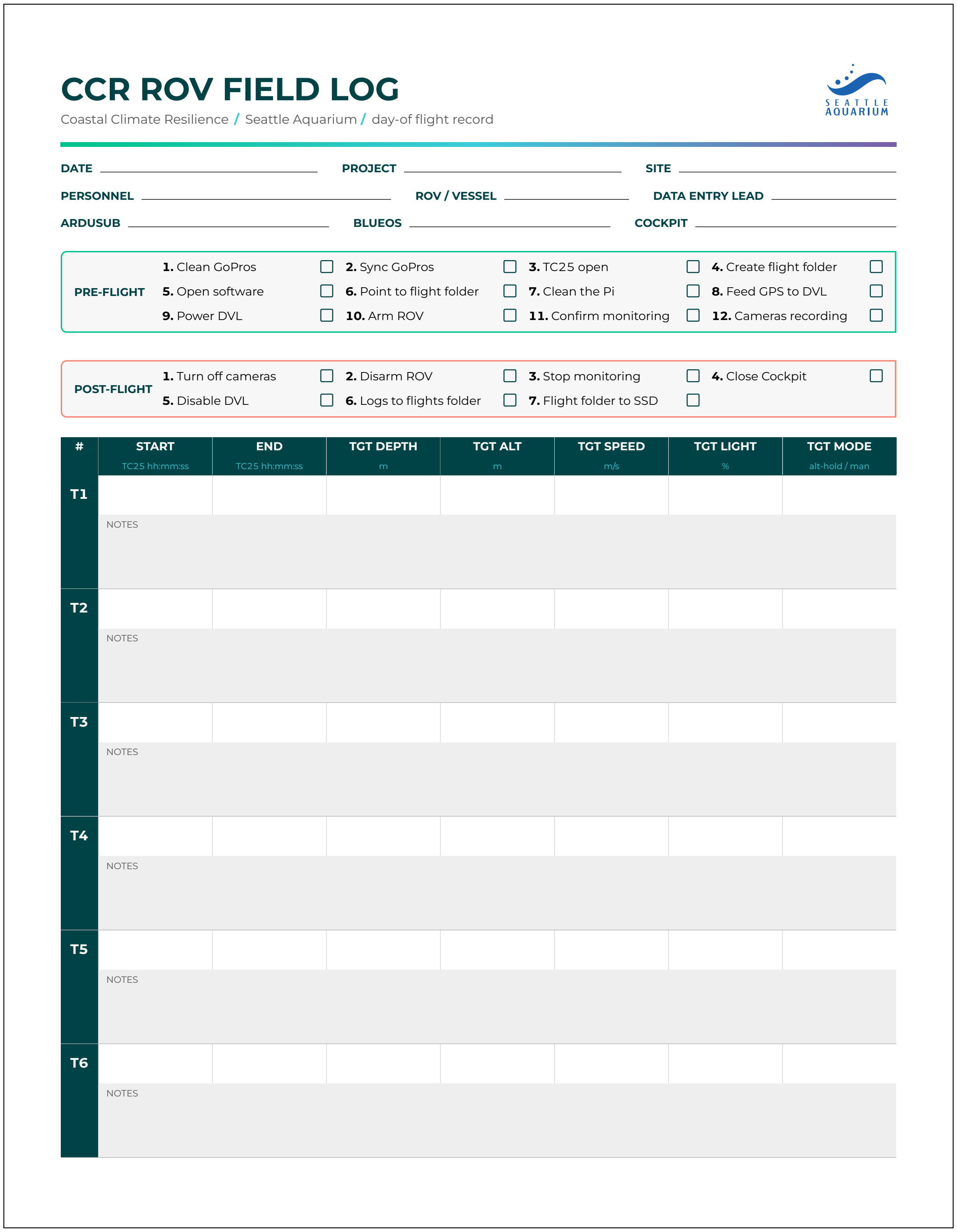

### CCR ROV FIELD LOG

Coastal Climate Resilience / Seattle Aquarium / day-of flight record

SEATTLE AQUARIUM

DATE ____________ PROJECT ____________ SITE ____________

PERSONNEL ____________ ROV / VESSEL ____________ DATA ENTRY LEAD ____________

ARDUSUB ____________ BLUEOS ____________ COCKPIT ____________

**PRE-FLIGHT**

| | | | |
|---|---|---|---|
| **1.** Clean GoPros ☐ | **2.** Sync GoPros ☐ | **3.** TC25 open ☐ | **4.** Create flight folder ☐ |
| **5.** Open software ☐ | **6.** Point to flight folder ☐ | **7.** Clean the Pi ☐ | **8.** Feed GPS to DVL ☐ |
| **9.** Power DVL ☐ | **10.** Arm ROV ☐ | **11.** Confirm monitoring ☐ | **12.** Cameras recording ☐ |

**POST-FLIGHT**

| | | | |
|---|---|---|---|
| **1.** Turn off cameras ☐ | **2.** Disarm ROV ☐ | **3.** Stop monitoring ☐ | **4.** Close Cockpit ☐ |
| **5.** Disable DVL ☐ | **6.** Logs to flights folder ☐ | **7.** Flight folder to SSD ☐ | |

| # | START TC25 hh:mm:ss | END TC25 hh:mm:ss | TGT DEPTH m | TGT ALT m | TGT SPEED m/s | TGT LIGHT % | TGT MODE alt-hold / man |
|---|---|---|---|---|---|---|---|
| T1 | | | | | | | |
| | NOTES | | | | | | |
| T2 | | | | | | | |
| | NOTES | | | | | | |
| T3 | | | | | | | |
| | NOTES | | | | | | |
| T4 | | | | | | | |
| | NOTES | | | | | | |
| T5 | | | | | | | |
| | NOTES | | | | | | |
| T6 | | | | | | | |
| | NOTES | | | | | | |

Figure S1: The one-page ROV field log, printed for each survey day: day and personnel metadata, software versions, pre- and post-flight checklists, and a record of each transect's TC 25 start and end times and target depth, altitude, speed, light level, and flight mode, with space for notes.

#### Field protocols

Here we provide complete protocols to (**1**) set up the ROV and topside command console, and (**2**) conduct ROV benthic survey flight operations, including within canopy-forming bull kelp forests.

> **Software:** Since 2026, `rov_flight_ops` and `rov_imagery_processing` (Section S4) automate several of the steps below — flight logging, network checks, transect-time entry, log download, flight reports, and imagery import. Where a step is supported, it is noted in the protocol and checklists.

##### Reference links

- BlueROV2 Software Setup: `https://bluerobotics.com/learn/bluerov2-software-setup/`
- WaterLinked DVL Networking: `https://docs.waterlinked.com/dvl/networking/`
- WaterLinked UGPS Network Settings: `https://docs.waterlinked.com/underwater-gps/network-settings/`
- WaterLinked BlueROV Integration (U1): `https://docs.waterlinked.com/underwater-gps/integration/bluerov-integration-u1/`
- Advanced Navigation GNSS Compass Quick Start: `https://docs.advancednavigation.com/gnss-compass/QuickStart.htm`
- GoPro Labs firmware: `https://community.gopro.com/s/article/GoPro-Labs`

##### Topside setup

> **Important — Do these first:** The WaterLinked G2 box and the U1 locator both require a GPS lock before use, which can take **up to 10 minutes**. Power them on as early as possible so they can acquire a lock while you complete the remaining steps.

###### Step 1: Anchor and initial power

1. Anchor the vessel securely above the ROV survey location.
2. Open the Pelican case and connect the internal power strip to vessel power.
3. Turn on the rugged laptop inside the Pelican case (pre-connected to the power strip).
4. Launch `rov_flight_ops` and choose the day's flight folder on the **Monitoring** tab. The flight recorder then starts automatically each time the ROV is armed.

###### Step 2: WaterLinked UGPS — turn on early

1. Power on the WaterLinked G2 topside box. It must obtain a GPS lock before the UGPS system will function; this can take up to 10 minutes.
2. Power on the U1 locator. It also requires a GPS lock. Set it aside in a clear-sky location while continuing setup.

###### Step 3: FXTI box connection

1. Connect the FXTI box to the laptop via USB.

> **Note:** If using the Outland Technology Power Supply, use the dedicated FXTI port on the right side of the Outland box instead.

**Step 4: WaterLinked antenna deployment**

1. Lower the rigid WaterLinked G2 antenna into the water until it is approximately **1 m below the surface**. Lock it into place.
2. Connect the antenna cable to the G2 box.
3. Connect the G2 box to the Ethernet switch inside the Pelican case.

**Step 5: Advanced Navigation GNSS compass**

1. Mount the GNSS compass on top of the vessel with an **unobstructed view of the sky**.
2. Connect a Power-over-Ethernet (PoE) cable to the compass and route it into the Pelican case.
3. Connect the PoE cable to the PoE switch inside the case (powered by the internal power strip).
4. Connect the PoE switch to the Ethernet switch.
5. Connect the Ethernet switch to the laptop's Ethernet port.
6. Confirm the Ethernet switch is also connected to the internal power strip.

**Step 6: Outland power system and ROV tether**

> **Important:** Do **not** power on the Outland system until the tether is connected to both the Outland topside unit **and** the ROV.

1. Connect the power cable and grounding cable from the Outland power system to the power strip.
2. Attach the ROV tether to the tether port on the Outland system.
3. Apply **Molykote 44 Medium High-Temperature Grease** to the wet-mateable tether connector pins on the aluminum enclosure.
4. Mate the tether connector to the ROV's aluminum enclosure.
5. Power on the Outland system. The ROV will now receive power.

**Step 7: Confirm ROV power-on sequence**

1. **First chime/beep set:** Indicates the power connection is established.
2. **Second chime/beep set** (approx. 1 minute later): Indicates the autopilot is active and thrusters are armed but not spinning.

**Network configuration**

The ROV connects to the laptop via USB, while the external Ethernet switch connects via the laptop's Ethernet port. These two interfaces must be joined using a **network bridge**; full instructions are available at https://docs.waterlinked.com/underwater-gps/integration/bluerov-integration-u1/.

Once the network bridge is active, verify each interface in a browser:

1. **BlueOS (ROV):** Navigate to `192.168.2.2`. Confirm the ROV is visible and connected.
2. **Advanced Navigation GNSS Compass:** Navigate to `http://gnss-compass.local/`
   - Default username: `admin`  Default password: `password`

   > **Note:** If `gnss-compass.local` does not resolve, locate the device IP via the router's connected-device list or a network scanner (e.g., `nmap -sn 192.168.2.0/24`).

3. **UGPS Topside:** Navigate to `192.168.2.94`
   - Click **System Status** (top left). All items should show green checkmarks *except* the locator — this is expected, as the locator is above water and not yet in range of the antenna.
   - **External compass note:** When configuring antenna placement, measure distances from the **GNSS compass** to the antenna, not from the G2 box.
4. **DVL:** Navigate to `192.168.194.95`. Confirm the DVL is connected and responsive.

   > **Overheating risk:** The DVL can overheat if left out of the water for an extended period. Disable the acoustics via the **Configuration** page in the DVL GUI while the ROV is topside. **Remember to re-enable the acoustics after deploying the ROV.**

> **Network check:** With the tether connected, `rov_flight_ops`'s **Check the network** (Monitoring tab) reports which adapter carries the tether, whether the network bridge is healthy, and whether Windows may power the adapter down; **Save to flight folder** keeps the result with the day's logs.

##### ROV prep

1. Connect the Xbox game controller to the laptop.

   > **Recommendation:** Use a wired (USB) connection rather than Bluetooth, which has been found to be less reliable in the field.

2. Open **Cockpit** (or QGroundControl). The live video feed from the ROV should appear automatically.
3. Arm the ROV and briefly test each thruster to confirm the connection is functioning correctly.

##### GoPro setup

###### Step 1: Firmware and time sync

1. Download and install **GoPro Labs firmware** on the camera (see reference links above).
2. Connect the laptop to the internet via a mobile hotspot (required for GoPro Labs Precision Date and Time to load).

3. Insert a charged battery into the GoPro and power it on.
4. Point the GoPro at the QR code generated by GoPro Labs Precision Date and Time (Local). A brief checkmark on the GoPro screen confirms the time has been synced.

**Why time sync matters:** Syncing the GoPro's clock to the laptop ensures image timestamps align with the ROV telemetry logs, which is essential for accurate post-processing and georeferencing.

##### Step 2: Create a custom GoPro Labs preset

Create a custom preset in GoPro Labs with the following settings:

| **Setting** | Value |
|---|---|
| **Mode** | Photo |
| **Interval** | 3 s |
| **Lens** | Wide |
| **Output** | RAW |
| **EV Compensation** | 0 |
| **White Balance** | Native |
| **ISO Min–Max** | 100–200 |
| **Sharpness** | Low |
| **Color** | Flat |
| **Shutter Speed** | 1/300 (`!MEXPX=300`) |

**Shutter speed override:** The GoPro Labs shutter speed (`!MEXPX=300`) only takes effect if the GoPro preset's shutter speed is set to **Auto**. A manually set shutter speed in the preset will override the Labs value.

**White balance:** Do **not** use Auto White Balance. Auto WB has caused severe overcorrection in the field, producing nearly unusable images and significantly complicating post-processing. **Native** white balance preserves maximum flexibility for color correction in post.

##### Step 3: Mount the GoPro

1. Attach the GoPro to the ROV with the lens pointing **straight down**.
2. Secure the mount tightly so the camera remains horizontal and does not shift during the survey.

**Pre-dive checklist**

| Item | Done? |
|---|---|
| Vessel anchored securely | ☐ |
| G2 box and U1 locator powered on (GPS lock acquired) | ☐ |
| All GUIs loading correctly (BlueOS, UGPS, DVL, Compass) | ☐ |
| Network bridge active | ☐ |
| `rov_flight_ops` flight folder chosen; monitoring active | ☐ |
| Navigation readiness checked; EKF origin set for DVL-only dives (`rov_flight_ops` Navigation tab) | ☐ |
| DVL acoustics disabled (re-enable after deployment) | ☐ |
| ROV thrusters tested | ☐ |
| GoPro time-synced with laptop | ☐ |
| GoPro custom preset applied (correct shutter + native WB) | ☐ |
| GoPro mounted pointing down and secured | ☐ |
| U1 locator attached to ROV vertically | ☐ |

**ROV flight operations**

**Step 1: Deploy the ROV**

1. Turn on the GoPro to begin capturing images, then lower the ROV into the water.
2. Once the ROV is at least **1 m underwater**, the U1 locator will come within range of the antenna and the ROV's position should appear in the UGPS GUI.
3. Re-enable DVL acoustics via the DVL GUI Configuration page.
4. Engage **Stabilize** flight mode and pilot the ROV to the desired survey start location at the surface.

   > **Tip — compass reference:** While the ROV is still visible at the surface, note the compass heading in Cockpit and visually confirm which direction the ROV is pointing. This gives a reliable heading reference before descending.

5. Descend to the desired survey start depth.

**Step 2: Engage surftrak and set altitude**

1. Once the ROV is approximately **1 m off the seafloor**, engage **surftrak** mode.
2. Adjust the target altitude to the desired survey height. We typically use **0.8 m**.
3. Confirm the ROV is holding the target altitude before proceeding.

> **surftrak2 behavior:** With `surftrak2.lua` installed, adjusting the target altitude will cause the ROV to return to the new set altitude automatically after any manual altitude change,

> rather than holding the adjusted position.

**Step 3: Begin the transect**

1. Orient the ROV to the desired compass heading.
2. Once altitude is locked and heading confirmed, **record the start time** from the GoPro Labs Precision Local Time page (TC 25 time). This timestamp will be used to trim telemetry logs and images to the transect.
3. Begin the transect, maintaining a speed of approximately **0.1–0.15 m/s**.

   > **Monitoring speed:** Survey speed can be monitored in the DVL GUI or added as a widget in Cockpit. In areas with minimal current, 20–30% thruster power is typically sufficient. Thruster power can be adjusted using the **D-pad** on the game controller.

4. At the end of the transect, **record the end time** (TC 25 time). Both start and end times are needed to isolate the telemetry files and GoPro images to the transect interval during post-processing.

> **Entering times and pauses:** After the flight, type the TC 25 start and end times into `rov_flight_ops`'s **Transects** tab and press **Save**, which writes them to the flight folder as `surveys.json`. If a transect was interrupted (e.g., the vehicle disarmed or the video glitched), also record the start and end of the interruption and add it as a **pause**; imagery from a pause is then kept out of the transect, and telemetry from it is flagged rather than deleted. **Preview transects** confirms, against the dive profile, that the transects were flown and recorded as written.

**ROV retrieval**

**Step 1: Exit the survey area**

1. Switch the ROV from SurfTrak to **Depth Hold** mode.
2. Rotate the ROV until you can locate the tether in the camera feed. The tether tender should pull in slack to make the tether easier for the pilot to locate.
3. Follow the tether path back toward the vessel, keeping the ROV relatively close to the seafloor.

> **Kelp bed navigation:** Exit the kelp bed by **retracing the same path used to enter**. Do not cut through the kelp in a new direction, as this risks entanglement. Have the tether tender pull in as much slack as possible to reduce loose tether that could snag on stipes.
>
> Keep the ROV **close to the seafloor** during the return — bull kelp stipes become more twisted and dense toward the surface, increasing entanglement risk at shallower depths. Once clear of the kelp bed, speed can be increased to return to the vessel.

**Freeing a snagged tether:** If the buoyant tether rises into a dense tangle of stipes near the surface on the return path, it can typically be cleared with a coordinated two-person technique: (**1**) the tether tender places tension on the tether by gently but firmly pulling, while simultaneously (**2**) the pilot moves the ROV laterally in a desired direction. This coordinated action (**3**) temporarily opens a gap in the kelp stipes, allowing the small and highly maneuverable vehicle to pass and clearing the tether from the bundle.

##### Step 2: Surface and recover

1. Bring the ROV to the surface near the vessel.
2. Switch the flight mode to **Manual** before recovery. This disables altitude-hold and stabilization, preventing thrusters from spinning unexpectedly during retrieval.
3. Recover the ROV aboard the vessel.
4. Turn off the GoPro.

##### Post-dive checklist

| **Item** | **Done?** |
|---|---|
| Transect start time (TC 25) recorded | ☐ |
| Transect end time (TC 25) recorded | ☐ |
| Transect times and any pauses saved and previewed (`rov_flight_ops` Transects tab) | ☐ |
| Vehicle logs downloaded and verified before power-off (`rov_flight_ops` BlueOS logs tab) | ☐ |
| Flight report generated (`rov_flight_ops` Flight summary tab) | ☐ |
| ROV recovered and powered off | ☐ |
| GoPro powered off | ☐ |
| Tether fully retrieved and coiled | ☐ |
| UGPS antenna retrieved from water | ☐ |
| All subsea equipment rinsed with freshwater | ☐ |
| All equipment stowed in Pelican case | ☐ |
| GoPro imagery imported with `rov_imagery_processing` before the SD card is cleared | ☐ |

#### ROV positioning system

Here we detail the positioning hardware, modifications, and configuration summarized in the main text (Methods and Results: Positioning). Figs. S3–S5 show the wiring and topside hardware, and Table S2 lists the associated software and BlueOS Extensions.

##### Underwater acoustic hardware

We used Water Linked's Underwater GPS (UGPS) G2 USBL system, mounting the battery-powered U1 locator vertically on the ROV and positioning the acoustic antenna 1 m beneath the sea surface. In its default configuration, the three foldable receivers of the easily deployed UGPS antenna were not sufficiently immobilized once in the water: as the anchored vessel bobbed on the surface, these receivers shifted position, introducing error into the acoustic calculations and the broader position estimate. The antenna mount itself — a ball mount designed to allow flexible deployment angles — also moved with vessel motion and introduced additional error. We addressed these issues by using stainless-steel hose clamps to completely rigidize the collapsible receivers and foldable antenna body, and by immobilizing the antenna mount with a PVC framework that slotted into a pot-puller/davit mount on the vessel (Fig. S4a, e). These changes made the system somewhat less convenient to store and transport, but improved acoustic stability and overall UGPS performance (main text Fig. 2).

##### Topside GNSS compass integration

On the topside half of the system, we suspected that the GPS receiver and compass housed within the UGPS G2 box were introducing noise into the integration of topside and acoustic data. For ROV position estimation, we therefore bypassed these sensors by mounting an Advanced Navigation GNSS Compass on the vessel cabin roof (Fig. S4c), which provides highly precise position and heading. Our BlueOS Extension `wl_ugps_external` (Table S2) receives the compass's NMEA GGA (position) and HDT (heading) messages, broadcast over UDP, and forwards them to the G2 box API. This approach retained the acoustic–topside integration of the UGPS system within the G2 box while replacing its weaker topside sensors; the G2 box's own GPS receiver was still used to obtain a satellite fix to synchronize time. When configuring antenna placement in the UGPS interface, offsets must be measured from the GNSS compass rather than from the G2 box (Section S2).

##### DVL and sensor fusion

We estimated ROV altitude with a BR Ping1D and, from 2023, a Water Linked DVL A50, which was retained across $V1$–$V4$ for the additional precision of its four acoustic beams (see Section S5 for its mounting). By default, the DVL A50 outputs a single altitude value derived from the four beams; we modified the Water Linked DVL BlueOS Extension to also write the raw beam data to the telemetry log, so that future flight control could anticipate terrain changes before they appear in the averaged altitude.

DVL and UGPS data were fused through ArduSub's EKF3, together with compass and inertial data from the Navigator Flight Controller. For improved DVL+UGPS fusion, the UGPS extension was run with `EXTRA_ARGS=--ignore_gps` (Table S2). The full navigational suite requires an Ethernet switch within the ROV enclosure for the DVL, a second Ethernet switch in the command console for the GNSS compass, UGPS G2 box, and laptop, and a Power-over-Ethernet switch to power the GNSS compass (Figs. S3 and S5); the laptop's USB tether interface and its Ethernet interface are joined by a network bridge (Section S2). The two EKF configurations we used — DVL dead reckoning and acoustic + DVL — and the ArduSub parameters each requires are encoded as navigation profiles in `rov_flight_ops`, which checks them against the vehicle before a dive (Section S4). For a DVL-only dive, the EKF must also be given a geographic origin before arming (we use the `ahrs-set-origin.lua` applet; Table S2); without one, the vehicle navigates normally in its local frame but logs no latitude or longitude.

##### `surftrak` altitude performance

We quantified how closely `surftrak` held its target altitude on all 2024–2025 survey transects for which both the vehicle's telemetry logs (`.tlog`) and the transect time windows were available, using `code/surftrak_performance.py`. For each second of each transect, the script takes the flight mode from the autopilot's own heartbeat, the mean rangefinder (DVL) altitude, and the target altitude that ArduSub reports while in `surftrak` (its `RFTarget` value), and scores only seconds flown in `surftrak` with a valid altitude reading. Error is the realized altitude minus that logged target, so it reflects the altitude the pilot actually requested rather than a nominal survey altitude.

Of 138 transects with telemetry, 124 included some `surftrak` time and 118 were flown in `surftrak` for at least half of their duration (median 100%), spanning 27 survey days and 20.8 h of 1 Hz telemetry; the altitude reading dropped out for only 15 s of that time. Per-transect median targets ranged from 0.65 to 1.05 m. Realized altitude was within ±0.10 m of the target for 84.7% of the time and within ±0.20 m for 96.2%, with no systematic bias (median error −0.003 m; mean absolute error 0.057 m), and performance was similar in both years (Table S1; Fig. S2). The median transect spent 99.1% of its `surftrak` time within ±0.20 m of the target. The lowest values came from a few transects on which the target had been raised to 0.9–1.0 m and the vehicle held 0.1–0.2 m below it for extended periods. Across all `surftrak` time in the logs, on or off transect (31.9 h), the share within ±0.20 m was 94.8%.

Table S1: `surftrak` altitude-holding performance on survey transects flown in `surftrak` for at least half of their duration, by year; survey days are calendar dates. Error is the ROV's rangefinder altitude minus the target altitude logged by the autopilot, at 1 Hz; the last two columns give the share of time within ±0.10 and ±0.20 m of the target. MAE, mean absolute error. Computed from the vehicle's telemetry logs with `code/surftrak_performance.py`, which also writes per-transect and per-survey-day results.

| **Period** | **Survey days** | **Transects** | **Time (h)** | **Altitude (m), median (IQR)** | **Median error (m)** | **MAE (m)** | **±0.10 m (%)** | **±0.20 m (%)** |
|---|---|---|---|---|---|---|---|---|
| 2024 | 12 | 50 | 8.4 | 0.70 (0.67–0.76) | +0.004 | 0.052 | 86.6 | 96.7 |
| 2025 | 15 | 68 | 12.5 | 0.77 (0.73–0.81) | −0.010 | 0.060 | 83.5 | 95.9 |
| **All** | **27** | **118** | **20.8** | **0.75 (0.70–0.80)** | **−0.003** | **0.057** | **84.7** | **96.2** |

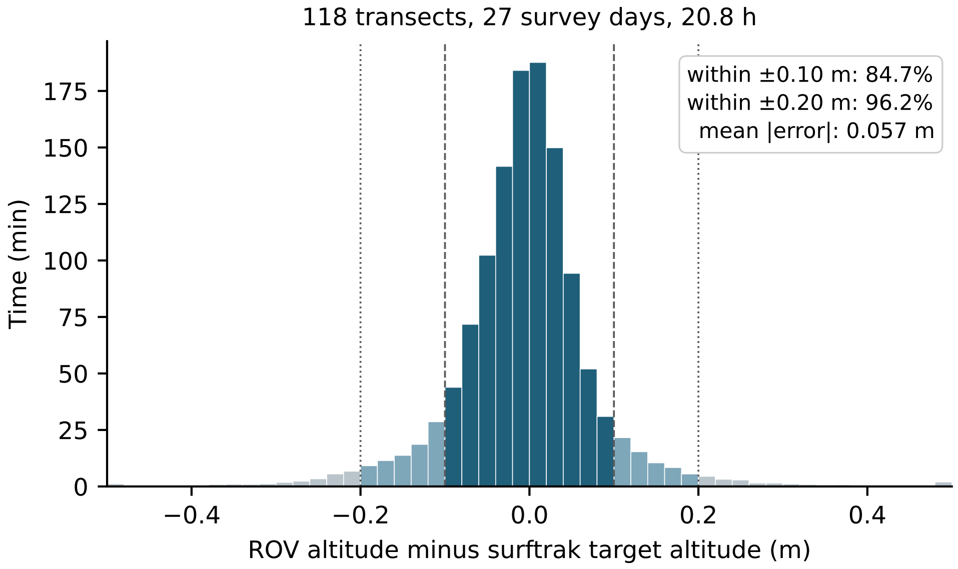

Figure S2: Distribution of `surftrak` altitude error — the ROV's rangefinder altitude minus the target altitude logged by the autopilot — for every second flown in `surftrak` on 118 survey transects across 27 survey days in 2024–2025 (20.8 h at 1 Hz). Dark bars fall within ±0.10 m of the target (dashed lines) and light-blue bars within ±0.20 m (dotted lines); the few errors beyond ±0.5 m are gathered into the end bins.

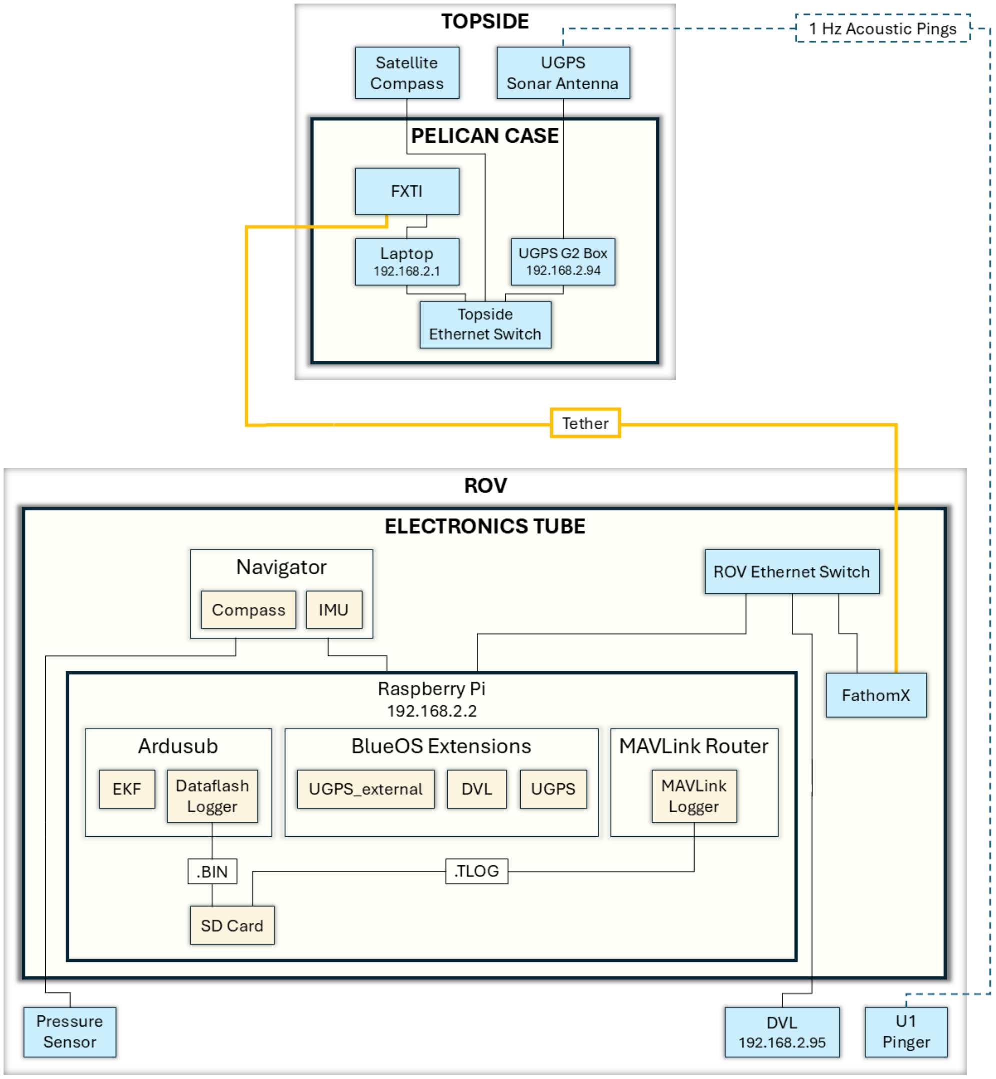

(a)

Figure S3: Conceptual wire diagram with the topside system including the full USBL positioning system and GNSS satellite compass, and the ROV electronics enclosure tube including the DVL.

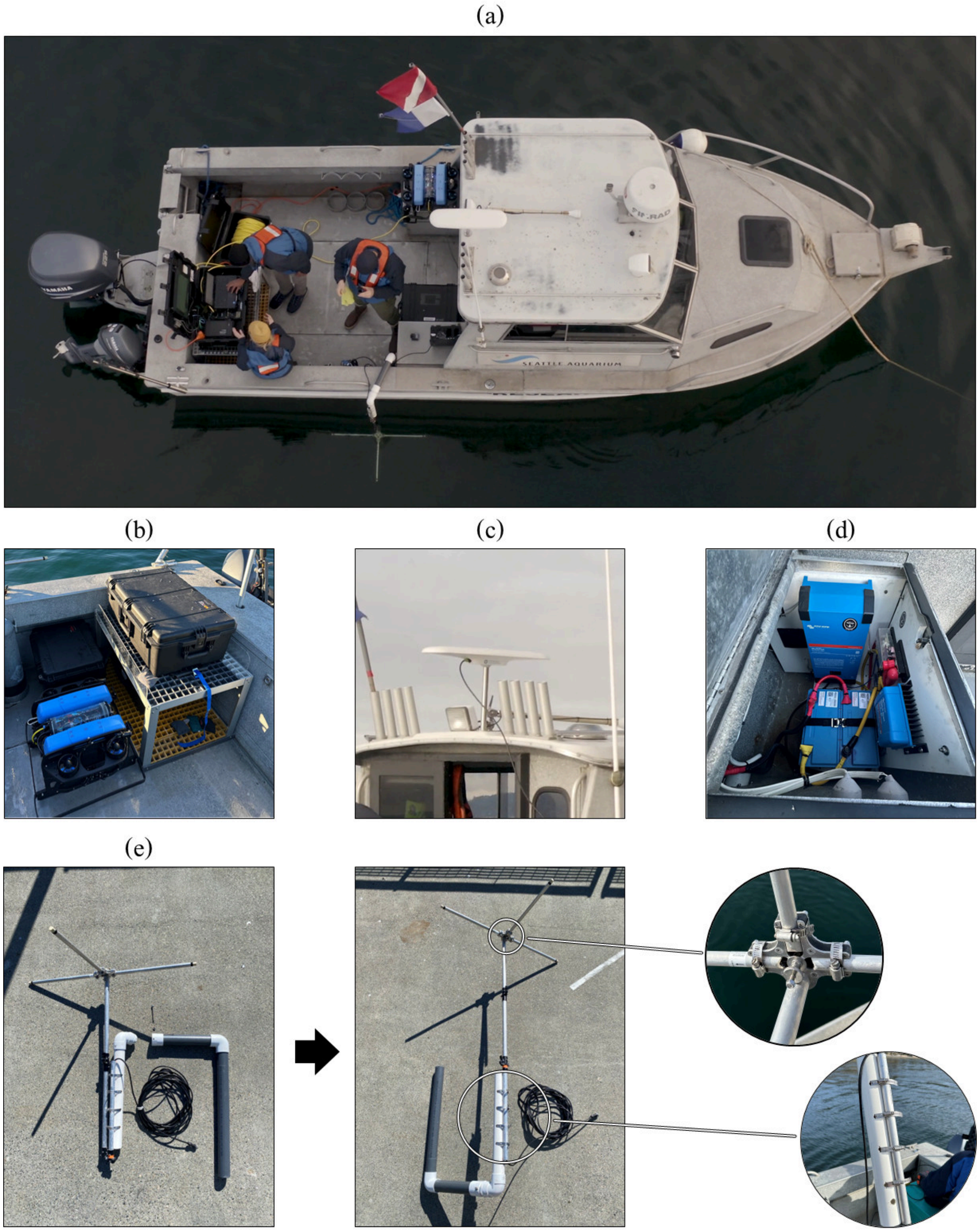

Figure S4: (a) Top view of the Seattle Aquarium vessel with command console by starboard transom. (b) The command console is strapped atop a Fiber Reinforced Polymer stand, with a second Pelican case tools and lights container stored underneath. (c) A GNSS satellite compass atop the vessel provides highly accurate topside GPS and compass information that is passed to the USBL system for integration with the acoustic data. (d) Two lithium-ion batteries and 2000 W inverter comprise the topside power supply for $V3$ and $V4$. (e) The revised WaterLinked UGPS antenna folded and disassembled for transport, and then extended and ready to deploy in the water. Note the inset stainless steel hose clamps that lock the otherwise movable antenna receivers and antenna body into place. The deployment of the UGPS antenna underwater can be seen in (a) above, with the PVC slotting into a pot-puller receptacle.

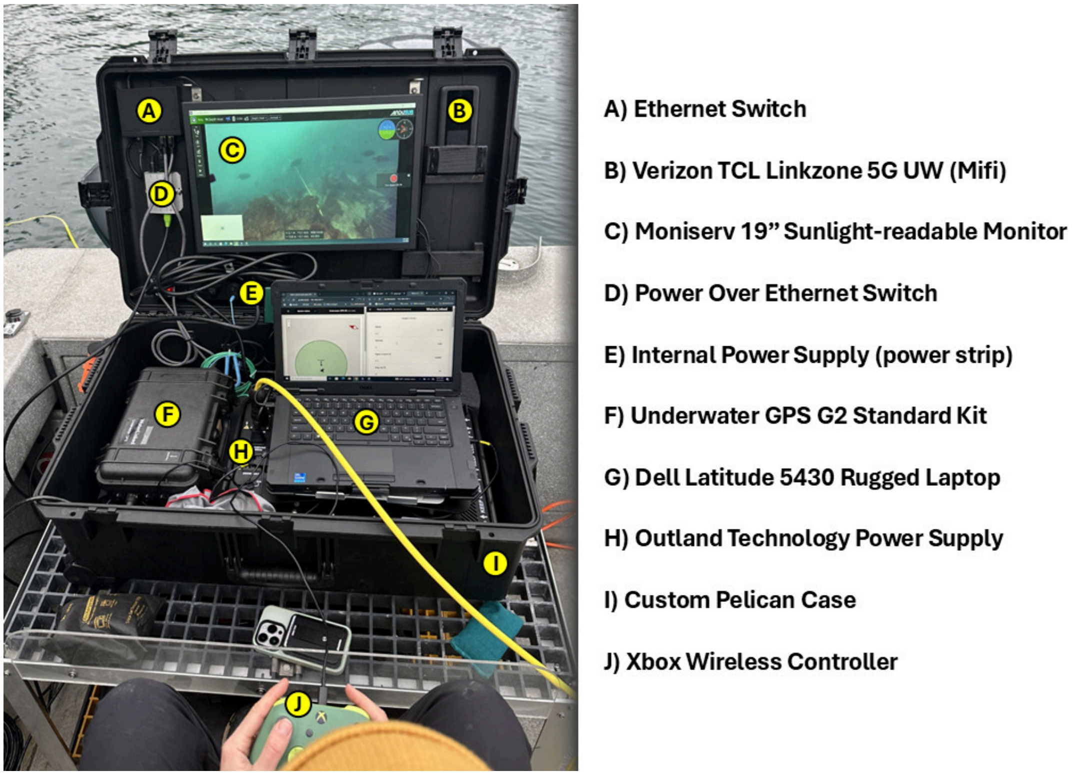

(a)

Figure S5: Our topside ROV command console and constituent components. The Outland Topside Power Supply system sits underneath the laptop, and the UGPS G2 box is affixed rigidly in place within the command console. The second screen enables multiple navigational display feeds, including the forward-facing ROV camera (upper screen), and the ROV position via the UGPS (left) and altitude and velocity information from the DVL (right) below on the laptop screen. All topside Ethernet switch and Power over Ethernet components are affixed and pre-connected within the command console.

#### Field operations and imagery processing software

We developed two open-source desktop applications, `rov_flight_ops` and `rov_imagery_processing`, that carry an ROV survey day from pre-dive checks to analysis-ready imagery (main text Methods and Results). Both are written in Python with a CustomTkinter interface, run on the Windows topside laptop, and are distributed in the CCR_ROV_survey_methods repository (folders `rov_flight_ops/` and `rov_imagery_processing/`). Each installs from a double-click launcher that builds a private Python environment with pinned, tested dependency versions and requires no administrator rights; thereafter it starts from a desktop shortcut and works without an internet connection.

The two applications are deliberately independent — neither imports code from the other — so that updating one cannot break the other. They share only data: the flight folder (Section S4), whose transect times (`surveys.json`) are written by `rov_flight_ops` and read by `rov_imagery_processing`, and a local cache of telemetry extracted from the vehicle's recordings, so that multi-gigabyte recordings are extracted once. Both keep a diagnostic log of every job, error, and interface stall on the laptop, and both are covered by automated test suites (768 tests for `rov_flight_ops`, 416 for `rov_imagery_processing`, and 157 for the transect extractor they share) that run on every change through continuous integration.

Before 2026, the same steps were run as stand-alone Python scripts (`tlog_to_csv.py`, `transect_map.py`, `rename_jpg_gpr_EXIF.py`, `optimize_path.py`, `meter_mark.py`), which produced the 2022–2025 data shown here and remain in the repository's `code/` folder. The applications grew out of a single program begun in August 2026, which we split into two standalone applications on 13 September 2026 to separate vehicle operations from imagery processing and to give each more room on the field laptop's screen. They were developed iteratively through the 2026 field season, with problems found on survey days fixed between surveys. Between 31 August and 18 September 2026, they (or their single-program predecessor) were used on all six Elliott Bay survey days (nine sites; 36 transects; 7,039 raw photographs sorted to transect), and `rov_flight_ops` recorded all six flights from 15–18 September (6.4 hours of topside telemetry; 31 GB of vehicle logs and imagery downloaded and checksum-verified), in addition to test flights at a nearshore test site. Development is ongoing; see the repository for current versions and documentation.

##### `rov_flight_ops`

`rov_flight_ops` runs on the topside laptop for the whole survey day. Its six tabs are numbered in the order a survey day uses them (Fig. S6), and a banner shows the vehicle's BlueOS, ArduSub, and Cockpit versions and whether the vehicle is connected and a flight is being recorded. Only two parts of the program can write to the vehicle — the navigation-profile and origin controls, and clearing old files — and an automated test enforces that every other request is read-only.

**1 Monitoring**

The flight folder is chosen first, before the dive (Fig. S6a). Pre-dive checks identify which network adapter carries the tether, whether the laptop's network bridge is healthy, and whether Windows may power the adapter down. A flight recorder then watches the vehicle and starts automatically when the ROV is armed, closing once it has been disarmed for 90 s so that a surface interval does not split one dive into several files. For each flight it records the topside laptop (processor, memory, storage, network, pilot-interface processes, and power) at 1 Hz, the tether's adapter counters at 10 Hz and pings at 5 Hz, and every autopilot parameter and software version at arming and disarming, noting any that changed. Live charts display these readings during the dive (Fig. S6b), and a recording failure (e.g., a full drive) is shown on every tab rather than passing silently.

**2 Navigation**
A live map of the ROV's track over an offline basemap, with the site origin and pilot-marked waypoints (Fig. S6c). Survey plans — measured lines, rotated rectangles, lane-filled grids, circles, and polygons, all dimensioned in meters — can be drawn and exported as GeoJSON, and a selected line provides live cross-track and along-track guidance. A navigation-readiness matrix reports, for each positioning source (USBL, vessel GNSS position and heading, DVL position and velocity, and ROV compass), whether it is *configured*, *receiving* new messages, *valid*, and *fused* by the EKF, showing a question mark where fusion cannot be established from what ArduSub publishes. Two navigation profiles (DVL dead reckoning; acoustic + DVL) check the vehicle's EKF parameters and explain why each requirement exists; applying a profile follows a deliberate review, apply, and read-back sequence that is locked by default and refused while the vehicle is armed. The page also shows whether an EKF origin has been applied, without which a DVL-only dive logs no geographic coordinates (Section S3). Recorded navigation sessions and transect CSVs can be replayed.

**3 Transects**
Transect start and end times are typed from the GoPro Labs precision timecode noted in the field (Section S2; Fig. S6d), together with any mid-transect pauses, and saved into the flight folder as `surveys.json`, which drives all downstream processing in both applications. Imagery taken during a pause is treated as off-transect, and telemetry rows from a pause are kept but flagged. A preview draws the dive profile with the transects marked (Fig. S6e), reading either the flight folder's recordings or — before anything has been downloaded — the autopilot's compact dataflash log directly from the vehicle, so that transects can be confirmed as flown and recorded before a camera card is cleared.

**4 BlueOS logs**
Lists the files on the vehicle's onboard computer — BlueOS `mcap` recordings, ArduSub dataflash (`.BIN`) logs, forward-camera video, imagery from an auxiliary MarineSitu C3 camera, and legacy `tlog` files — with the time span each recorded (Fig. S6f) and the transects each covers. Files are selected by type and period (this flight, today, transects only, or by hand), downloaded into the flight folder, and verified by size and SHA-256 checksum in a download manifest. Old files can then be cleared from the vehicle, guarded by a confirmed, current disarmed state, file metadata re-read immediately before deletion, and a deletion record written before anything is deleted.

**5 Flight summary**
Builds a flight report (PDF) from the day's logs. The report matches every recording to why it ended (e.g., an autopilot ground-station failsafe), places the tether-link state beside each disarm, compares pilot interfaces when more than one was used, and summarizes EKF, compass, GPS, DVL, and altitude-source health; topside resource use; parameter changes; and onboard-computer temperature. This tab also checks recording health, distinguishing a recording the vehicle never closed, which is repaired into a new copy, from a dive during which the vehicle stopped sending telemetry, and can substitute the dataflash log for telemetry when a recording failed.

**6 Analyze transects**
Produces per-transect 1 Hz CSVs — time; position from GPS, EKF, and DVL sources; tide-corrected depth; altitude; heading; velocity; distance; estimated image footprint and area; power;

light level; and survey state — and an interactive map, using the transect extractor mcap_to_csv, together with a sensor-health report. When a dive has no absolute position fix, the DVL track can be georeferenced from the origin stored in the vehicle's parameters, with a warning that the result is dead reckoning.

##### rov_imagery_processing

rov_imagery_processing runs on any laptop with the flight folder, in the field or the lab, and its four tabs follow the imagery from camera to analysis-ready files.

**1 Flight & transects**
Reads the transect times and pauses from surveys.json (or accepts them typed directly) and previews them against the dive profile, a check worth making before a camera card is cleared.

**2 Import photos**
Copies GoPro raw (GPR) photographs and JPEG previews directly from the SD card, which is never written to, into per-transect folders, keeping only frames taken during a transect; off-transect frames can optionally be kept separately. Each raw and preview pair is renamed to a shared capture-time name (YYYY_MM_DD_hh-mm-ss) taken from the preview's EXIF, replacing the earlier rename_jpg_gpr_EXIF.py and meter_mark.py scripts. JPEG previews can be stamped with a telemetry banner added above the frame, leaving every original pixel intact, and renamed with the values that decide whether a frame is usable (e.g., altitude, speed, and light power). Optionally, one photograph is selected per user-defined distance along each transect (e.g., every 1 m), with no photograph used twice and no distance accrued during pauses, producing a marks CSV and a track plot. Distance is integrated at sensor rate from the best available source — integrated EKF velocity, DVL odometry, GPS-derived velocity, or EKF position — each scaled by a calibration factor measured against 100 m tapes. On passes flown along the tapes on ten diver–ROV survey days in 2024–2025, EKF velocity recovered a median 99.3% (28 passes) and DVL odometry 96.3% (33 passes) of tape length, within 0.3% of the factors the program applies; the analysis script and per-pass results are in the repository (distance_calibration/). Integrating EKF velocity instead at the 1 Hz resolution of the transect CSVs moved 29% of one-meter marks to a different photograph. Each transect is also checked for agreement among distance sources and for straightness.

**3 Process photos**
Develops GPR raws through Adobe Lightroom Classic, using a bundled plug-in and a private scratch catalog, applying the standardized edits described in Section S6 — a centered $4606 \times 4030$ crop, chromatic-aberration removal, and AI Denoise at 50% — and exporting 16-bit ProPhoto RGB TIFFs. A preflight check precedes each batch, and progress through Lightroom's denoising and export is monitored and reported.

**4 Video**
Trims 4K GoPro video to each transect by stream copy (without re-encoding), cutting out pauses; builds telemetry composites at 4K, 1080p, or 720p that overlay the ROV's forward-camera video, gauges, and a telemetry panel on the downward-facing footage, with a matching 1 Hz CSV; and cuts short clips or places two videos side by side or stacked for comparison. Before compositing, the GoPro timecode is checked against the vehicle clock in two independent ways: the downward-facing footage's brightness against the ROV's light-power changes, and the rotation of the downward-facing image against the autopilot's yaw rate. A measured offset is applied only when the correlation is unambiguous and both halves of the transect agree; a transect flown dead

straight is reported as unverified rather than guessed. The ROV camera stream, which arrives over the tether at a variable frame rate, is re-timed from the recorder's capture timestamps and resampled so that the inset stays frame-accurate across long dives.

##### The flight folder

Every file from a survey flight is filed into one folder per site and day, so that the day's data can be backed up, shared, and reprocessed as a unit:

```
2026_09_17_EBM_W/
    surveys.json             transect times and pauses (Transects tab)
    logs/
        mcap/  BIN/  tlog/     recordings downloaded from the vehicle
        mcap_video/            forward-camera video extracted by BlueOS
        laptop_monitor_*.csv   topside laptop, 1 Hz          } written by
        network_fast_*.csv     tether adapter counters, 10 Hz } the flight
        network_pings_*.csv    tether pings, 5 Hz             } recorder
        flight_*.json          parameters, versions, and what changed
        nav_*.json, .jsonl     navigation session
        pi_downloads.jsonl     every download: source, size, time, SHA-256
        flight_report_*.pdf    flight report (Flight summary tab)
    photos/
        transects/T1/          GPR/  JPG_preview/  JPG_edited/  marks
        off_transect/
    transects/               per-transect CSVs and map
    videos/
        transects/T1/          4K trims
        composites/  clips/
```

##### Lessons from field testing

Field use in 2026 surfaced several failure modes that are silent without dedicated tooling, each of which the applications now detect or prevent:

- **Tether dropouts and failsafe disarms.** On 17 September 2026, the vehicle disarmed five times on ground-station failsafes. The flight report showed that the tether link had fallen silent five times (21 s in total; longest 5 s), that four of the failsafes followed 1–2 s of link silence, and that the link was held 96% of the time under Cockpit versus 100% under QGroundControl, with four failsafes versus one. This diagnosis, produced within minutes on the boat, had previously taken an evening of manual log review.
- **Telemetry that stops while the recording continues.** On 1 September 2026, the vehicle's MAVLink router stopped at 09:40:59 while the onboard recorder continued for another 57 minutes of video, leaving three of four transects with no telemetry in a structurally valid file. The recording-health check now distinguishes this from a file the recorder never closed, which can be repaired.
- **No EKF origin.** DVL-only dives on 17–18 September 2026 navigated normally but logged no geographic coordinates: origin coordinates had been entered as vehicle parameters before arming, but the EKF never applied them. The readiness matrix now shows whether an origin has been applied before the dive, and such dives can be georeferenced post hoc from the recorded origin parameters.
- **Clock offsets between GoPro and vehicle.** On a 14 September 2026 test flight, the GoPro timecode was 34.8 s behind the vehicle clock — undetectable by the light-ramp check on a

transect flown with steady lights, but measured from the vehicle's turns by the yaw-rate check.

- **Logs that grow while the vehicle is powered.** ArduSub appends to its dataflash log for as long as the vehicle has power, so every such download initially failed a strict size check; downloads that arrive longer than listed are now verified against the vehicle rather than rejected, whereas short (truncated) downloads still fail.

##### Limitations and ongoing development

Both applications currently run only on Windows, and automated raw development requires a licensed copy of Adobe Lightroom Classic. Several capabilities were written against documented BlueOS behavior and simulated vehicles in automated tests and are still being confirmed across vehicles and BlueOS releases, and extraction for the transect preview currently processes the vehicle's video stream as well as its telemetry, which costs disk time the preview does not need. Because ArduSub publishes its aiding mode rather than which individual sensors the EKF is fusing, the readiness matrix cannot always establish fusion and says so rather than guessing. We continue to develop both applications and welcome issues and contributions through the repository.

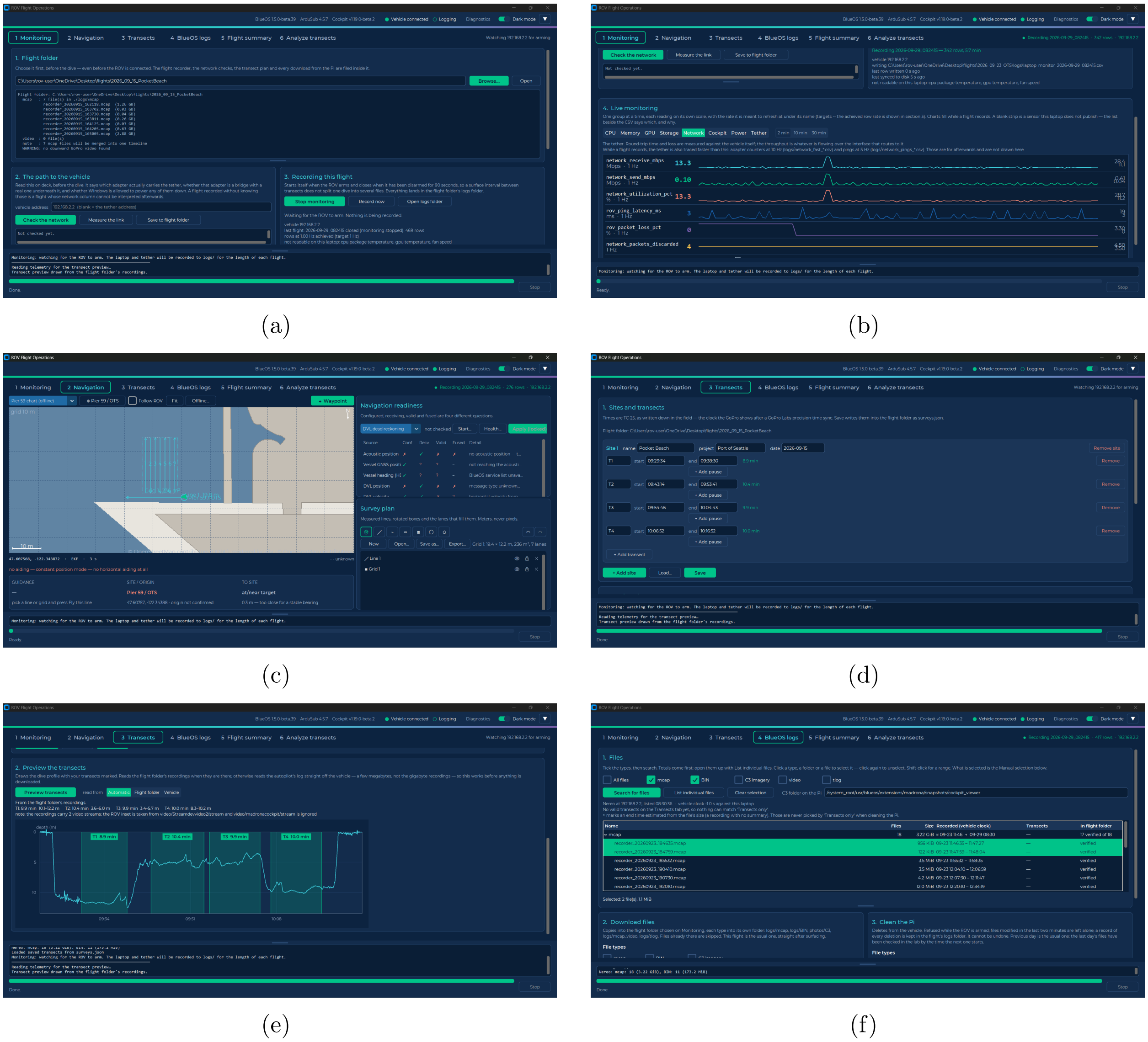

Figure S6: `rov_flight_ops` in use during 2026 field operations (Section S4). (a) Monitoring tab with the flight folder for a 15 September 2026 Elliott Bay survey selected, listing its seven vehicle recordings, while the flight recorder waits for the ROV to arm. (b) Live monitoring of the topside laptop and tether (network readings shown) while a flight records. (c) Navigation tab showing the offline basemap with a planned survey line and lane-filled survey grid, the navigation-readiness matrix, and the survey-plan tools. (d) Transects tab with the four 15 September transects entered as start and end times. (e) Transect preview: the dive profile (depth against local time) with the four transects marked, drawn from the flight folder's recordings. (f) BlueOS logs tab listing the recordings on the vehicle's onboard computer, with their recorded spans and whether each is already in the flight folder.

Table S2: Software and firmware used for ROV survey operations. All BlueOS extensions are installed and managed through the BlueOS web interface (`192.168.2.2`). Version numbers reflect those in use at the time of writing. See the CCR_ROV_survey_methods repository for current versions.

| Name | Type | Version | Description |
|---|---|---|---|
| BlueOS | software | v1.4.3-beta.7 | *Web-based platform for managing the ROV from its onboard Raspberry Pi computer; hosts all BlueOS extensions below.* |
| ArduSub | firmware | v4.5.0 (stable) | *Autopilot firmware running on the ROV flight controller; handles flight modes and sensor fusion via the EKF3 filter.* |
| Cockpit | software | v1.17.0 | *Browser-based ROV pilot interface; displays live video feed, telemetry widgets (depth, altitude, heading, speed), and controller input.* |
| BlueOS-Water-Linked-DVL | BlueOS extension | v1.0.8 | *Integrates the WaterLinked DVL A50 with the BlueROV2; provides velocity and altitude measurements to ArduSub for flight modes including `surftrak` and dead-reckoning navigation.* |
| blueos-ugps-extension | BlueOS extension | v1.0.7 | *Integrates the WaterLinked UGPS system with ArduSub by translating acoustic position data into MAVLink `GPS_INPUT` messages. Run with `EXTRA_ARGS=--ignore_gps` for improved DVL+UGPS sensor fusion.* |
| wl_ugps_external | BlueOS extension | v1.1.0-beta.1 | *Receives NMEA GGA (position) and HDT (heading) messages broadcast via UDP from the Advanced Navigation GNSS Compass, and forwards them to the WaterLinked UGPS G2 box API. This is the pathway by which topside vessel position and heading are fed into the UGPS system, replacing the G2 box's internal GPS with the higher-accuracy satellite compass.* |
| surftrak_fixit | BlueOS extension | v1.0.0-beta.2 | *Diagnoses and resolves issues with the `surftrak` altitude-hold flight mode.* |
| surftrak2.lua | ArduSub Lua script | — | *Modifies `surftrak` behavior so that manually adjusting the target altitude causes the autopilot to re-engage at the new set point, rather than holding the adjusted position indefinitely.* |

*continued on next page*

*Table S2 continued*

| Name | Type | Version | Description |
|---|---|---|---|
| ahrs-set-origin.lua | ArduSub Lua script | — | *Applies a stored latitude/longitude as the EKF origin before arming, which a DVL-only dive needs in order to log geographic coordinates (Section S3).* |
| `rov_flight_ops` | desktop application | Sep. 2026 | *Topside field-operations application: flight folder and automatic flight recording, live navigation map and sensor readiness, transect times and preview, verified log download from the vehicle, flight reports, and per-transect CSVs (Section S4).* |
| `rov_imagery_processing` | desktop application | Sep. 2026 | *Imagery application: per-transect photo import and distance-based photo selection, batch raw development in Adobe Lightroom Classic, and video trimming and telemetry compositing (Section S4).* |

#### Custom ROV hardware

##### Payload skid modifications, kelp guards

All iterations of the ROV utilized minor modifications to the ROV's primary frame ($V1$) or lower BR Payload Skid ($V2$–$V4$) to enable attachment of GoPro cameras and the Doppler Velocity Log (DVL). Unless stated otherwise, we utilized $\frac{1}{4}$ inch 316 stainless steel bolts and lock-washers/lock-nuts. Our $\frac{1}{4}$ inch hardware worked well, though it may be slightly excessive — a slightly smaller, lighter-weight hardware solution would likely be sufficient.

We cut away part of the BR Payload Skid with a jig saw to enable two GoPros to be mounted with an unobstructed downward-facing view. We first mounted the Ping1D towards the front and in the middle of the Payload Skid, though relatively quickly in 2023 upgraded to the Doppler Velocity Log and placed it in this same location (see Fig. S7 a, c, respectively, for $V1$ and $V4$ views of the ROV's payload skid). Apart from the lower Payload Skid, two small forward-facing GoPro mounts were installed adjacent to the four inch main electronics enclosure dome to enable GoPro cameras to be mounted facing forward.

Two 22 x 2 inch pieces of HDPE were bolted to the inside frame to provide attachment points for our various $V1$–$V4$ iterations of benthic illumination and our kelp guards. To prevent anything from getting caught within the lights that protruded out horizontally from the ROV's body — particularly stipitate kelp such as bull kelp stipes (the vertical structure bull kelp forms in the water) — we created a simple framework of HDPE that surrounded $V1$–$V3$'s lights. A 1 inch thick wood forming block was cut in the desired kelp guard shape, and a handheld heat gun was used to heat either a 1.5 or 0.75 inch wide, 36 inch length, and 0.25 inch length HDPE. The now-flexible HDPE was bent around the desired shape, and screws were temporarily installed into the HDPE and forming block to hold the bent shape. A towel with cold water was held over the bent joints to further solidify the HDPE's new shape. These two newly-shaped kelp guards were bolted onto the aforementioned two inch framework, with fore and aft connection between the two halves provided by two small BR metal rods (`MISC-BR - BR-100025 BROV2 Payload Skid Standoff, AL6061-T6`). These two horizontal rods prevented vertical material from being entrapped in the center and rear of the vehicle, while still enabling the built-in ROV camera to have an unobstructed view. See Fig. S7 a for an early $V1$ version of these kelp guards. By 2025 we reduced their thickness and footprint to minimize drag and nestle minimally around the $V2$ and $V3$ Kraken lights (see main text Fig. 1 b).

##### Light mounts

All $V1$–$V4$ lights were mounted laterally, off the left and right sides of the frame, and flush with the front and rear of the vehicle, maximizing their distance from the downward-facing camera; see, e.g., Fig. S7 a and c for the underside view of $V1$ and $V4$, respectively. To provide this custom spacing and placement for the benthic-illumination across $V1$–$V3$, we utilized the following aluminum underwater photography mounts:

- Ultralight Ball and $\frac{1}{4}$-20 threaded hole, female (url link).
- Ultralight New Style Clamp with $\frac{1}{4}$-28 thread (url link).
- Ultralight Aquavision Type Mount-Narrow (url link).

For $V1$, we used a Dremel rotary tool to cut away a small portion of aluminum from the third item above, enabling a small piece of HDPE to be attached as an adapter via which we bolted in place the BR Lumen Lights (Fig. S7 a). The same aluminum mounts were utilized for $V2$ and $V3$, however the weight of the Kraken batteries and lights caused the mounts to sag and twist under load, particularly the base mount, which would rotate around its single $\frac{1}{4}$ inch bolt. To affix the mounts and Kraken lights in place, we tapped two screw holes into the base aluminum mount, and inserted small screws into the HDPE, thereby preventing twisting of the base mount (Fig. S7 b). These mounts were originally intended to provide both freedom of motion to take advantage of variable angles, and the ability to lock the lights in place. In practice, the integrated BR Lumen Lights ($V1$) and battery powered Kraken lights ($V2$, $V3$) — due to their very wide beam angles (135° and 125° degrees, respectively) — adjusting the light angle was of limited utility, and we typically ran them facing straight down. Therefore, for any entities seeking to replicate $V1$–$V3$, we recommend spacing the lights out and away from the downward-facing camera akin to the methods described here, as that placement helped reduce back-scatter and improve image quality. However, there are mounting options available that are fixed and avoid the adjustable angle aspect of our mounts, which minimizes complexity and would eliminate undesired light angle movement/error. If custom angles are desired, a fixed 3D printed design that captures the desired light angles would be a better approach.

To mount our $V4$ integrated DeepSea lights we experimented with two methods, the first being default L-brackets which provided very stable mounting (Fig. S7 c). Second, we developed a custom aluminum bracket that enabled variable angles to be easily fabricated via https://sendcutsend.com/. The 3D design for these custom mounts can be viewed here at Onshape, and the mounts can be seen on ROV *Lutris* in Fig. 1 c. As the DeepSea lights had a much tighter 75° beam angle, we developed two interactive Google Colab notebooks to simulate ROV height above the seafloor, light placement, and beam angle to explore how adjusting the lateral/forward angles of the lights could be adjusted to maximize the benthic light's field of view coverage.

Both are in the repository (`lighting/beam_visualizers/`) and run in Google Colab or any Jupyter environment:

- 2D forward-facing view including percent-cover lumen bands for the DeepSea lights: ROV_Beam_Visualizer_LuxBands.ipynb
- 3D view with cones used to simulate the beam angle and light coverage from the four DeepSea lights: 3D_ROV_Beam_Visualizer.ipynb

These simulations were used to support the use of the narrower, 75° beam angle associated with the DeepSea lights, which made more efficient use of lumens, and the use of 10° lateral and forward offsets enabling a slightly wider illuminated field of view, while still ensuring some beam overlap to handle instances of variable terrain.

##### Integrated lights

We developed a custom ROV lighting control subsystem to provide real-time, pilot-adjustable illumination for benthic imaging surveys. The system uses four externally mounted SeaLite / Kraken Solar Flare Mini LED lights controlled by a Raspberry Pi Pico 2 housed in a Blue Robotics 1 L watertight enclosure. The enclosure contains the microcontroller, RS-485 transceiver, voltage conversion hardware, terminal blocks, WetLink penetrators, and an INA-series current / voltage monitor for power measurements. Lighting intensity is commanded from the BlueROV through an auxiliary servo PWM output configured in BlueOS and mapped to a joystick axis or button in Cockpit / QGroundControl. The Pico reads the incoming PWM signal, maps pulse width to 0–100% brightness in increments of 10%, and sends commands to the lights over UART using the DSPL SeaSense protocol. The firmware includes filtering, a brightness deadband, power-monitoring support, and a fail-safe that turns the lights off if the PWM signal is lost. This design provides an independently housed, modular lighting-control architecture that can be adjusted during flight while preserving compatibility with the existing BlueROV control workflow.

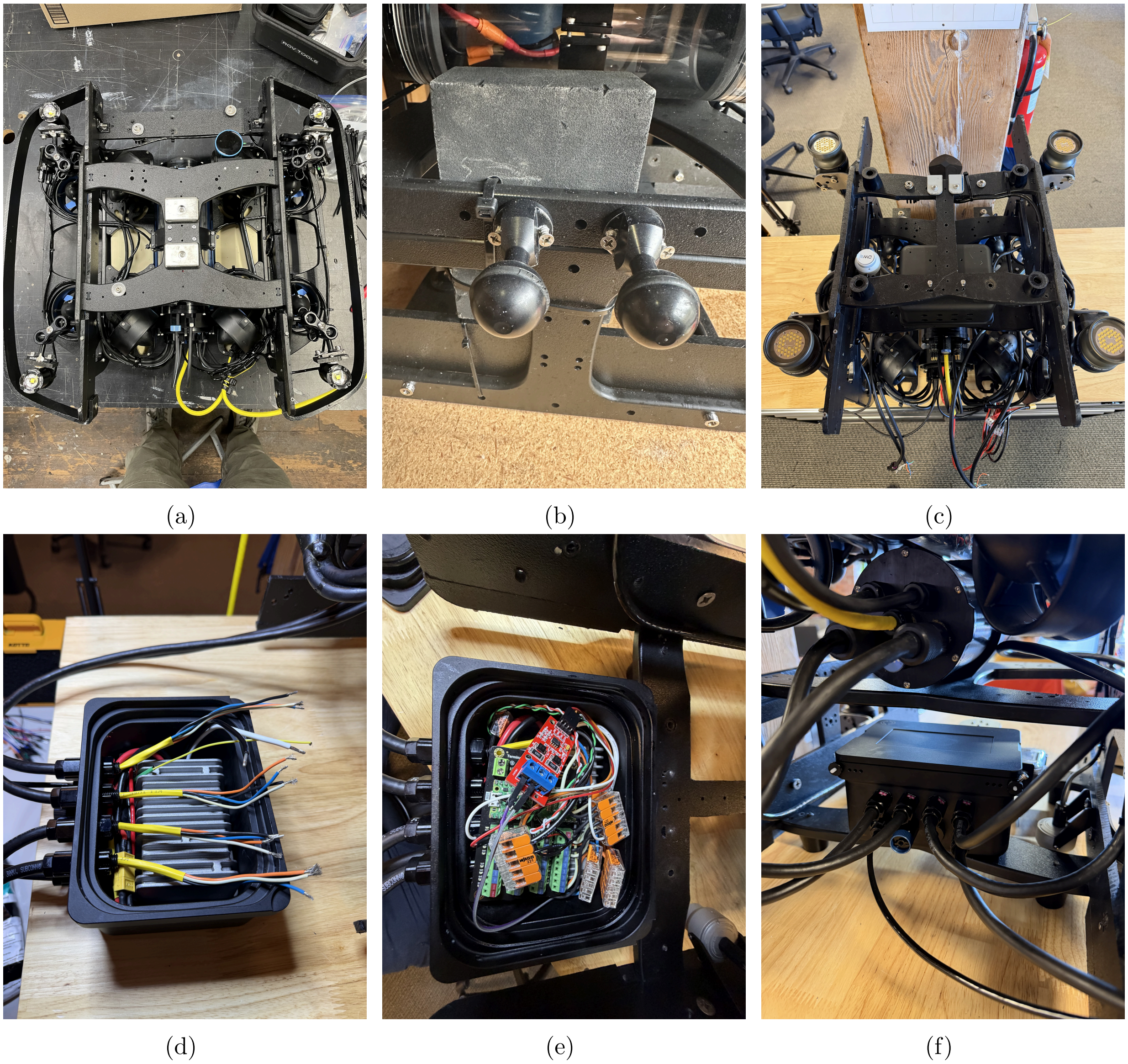

Figure S7: (a) $V1$ configuration with integrated BR lights and the aluminum lights mounts. (b) aluminum light mounts adjusted for the heavy $V2$ and $V3$ battery-powered Kraken lights, and tapped for stainless steel screws to prevent any rotation from the mount under the weight of the Kraken lights. (c) An initial version of the $V4$ integrated lights before we switched to the custom designed aluminum mounts (see Fig. 1 c in the main text for these custom designed mounts). (d) The 1 L enclosure for the $V4$ integrated lights with the voltage boost converter and conductor terminations for the four DeepSea lights. (e) The full complement of the 1 L enclosure Raspberry Pi Pico, RS485 converter, and WAGU connectors enabling rapid wiring and prototyping. (f) The completed 1 L enclosure mounted to the rear of the $V4$ payload skid.

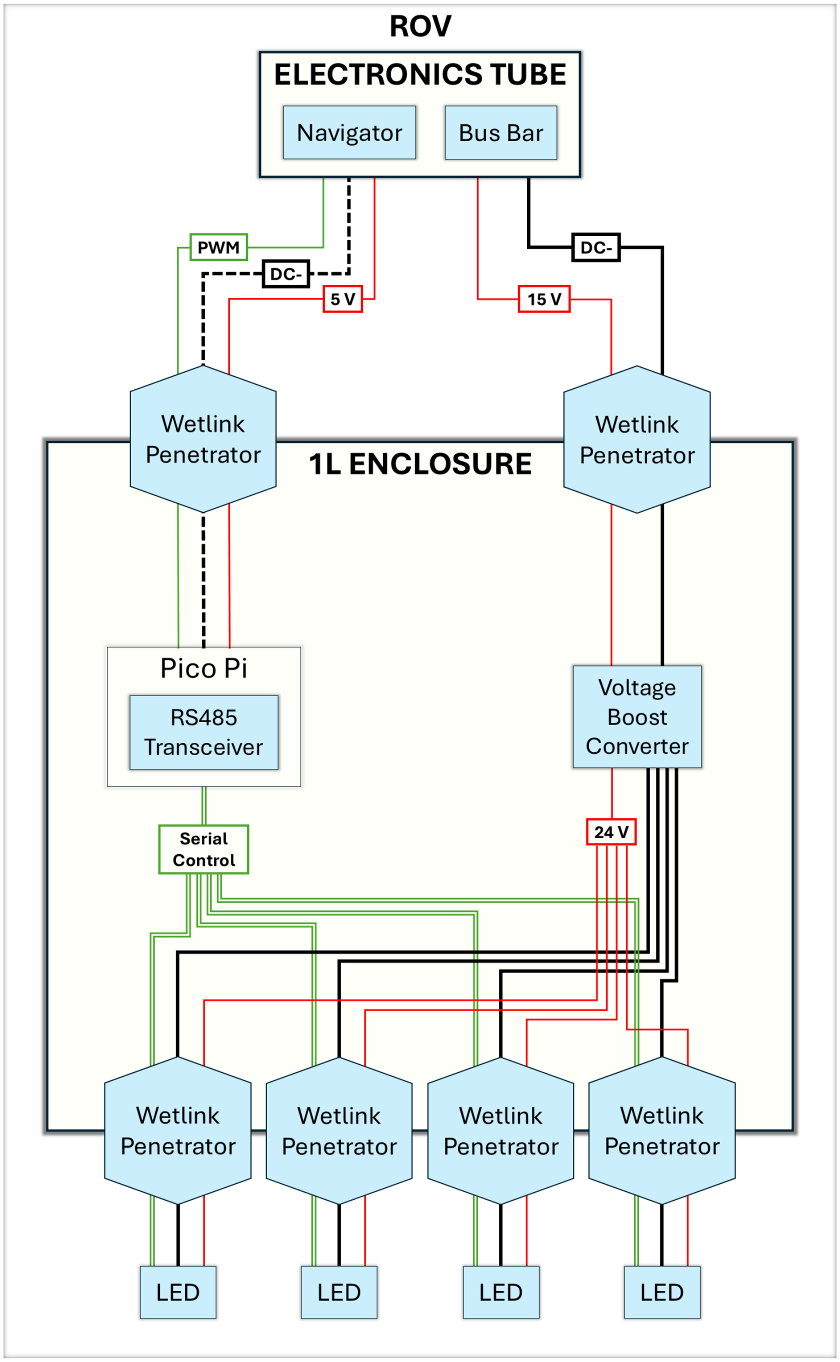

(a)

Figure S8: Wire diagram for our customized lighting control housing contained within a 1 L BR enclosure. Not shown are the single BR blanking plug and the pressure relief valve. The two DC- black lines (solid and dashed) represent the potential for two different ground planes due to power emanating from two different locations, the 15 V ROV busbar (solid black) and 5 V Navigator Flight Controller (dashed black).

Table S3: Bill of materials for the custom ROV lighting-control enclosure. Prices retrieved September 29th 2026 and do not include sales tax, shipping, or tariff surcharges. The reported total cost for the listed components is $662.89.

| Item | Qty. | MFN part no. | Unit cost | Description |
|---|---|---|---|---|
| Enclosure 1 Liter Box | 1 | BR-102603 | $255 | *1 Liter watertight box.* |
| WetLink Penetrator – M10 8.5 mm LC | 6 | BR-100870-085 | $13 | *Four WetLink penetrators for the four DeepSea lights, plus two WetLink penetrators for the power cable carrying 15 V from the main electronics tube.* |
| WetLink Penetrator – M10 4.5 mm LC | 2 | BR-100870-045 | $13 | *Two WetLink penetrators for the PWM cable connected to the Navigator in the main electronics tube.* |
| Pressure Relief Valve | 1 | BR-100804 | $32 | *Pressure relief valve for the lighting-control enclosure.* |
| Hole Plug M10 blank | 1 | BR-100434-010 | $6 | *Plug for the unused hole in the 1 Liter Box.* |
| Lumen/Gripper cable, 3 conductors, 22 AWG | 2 | BR-101050 | $16 | *Cabling for the PWM signal; paired with M10 4.5 mm LC WetLink penetrators.* |
| High Power Cable, 2 conductors, 12 AWG | 2 | BR-101093 | $19 | *Cabling for 15 V power from the main electronics tube; paired with M10 8.5 mm LC WetLink penetrators.* |
| Voltage Booster Converter | 1 | ATOWG-12S2420 | $144.49 | *12–24 V DC–DC boost converter. The project bill of materials notes that the 480 W version should be selected.* |
| Microcontroller, Raspberry Pi Pico 2 | 1 | SC1631 | $6.12 | *Microcontroller used to read the ROV servo PWM signal and command light intensity.* |
| Microcontroller Connection Board & Converter | 1 | DFR0924 | $19.90 | *Carrier board that provides voltage conversion for the Pico and screw-terminal connections for easier wiring.* |
| Current Monitor | 1 | — | $12.88 | *INA226 current and voltage monitor used for lighting-system power measurements.* |

*continued on next page*

*Table S3 continued*

| **Item** | **Qty.** | **MFN part no.** | **Unit cost** | **Description** |
| --- | --- | --- | --- | --- |
| Terminal Blocks | 1 | — | — | *Terminal blocks or WAGO connectors used to connect the power lines from the LEDs to the output of the converter.* |
| RS-485 Transceiver | 1 | BOB-10124 | $12.50 | *RS-485 transceiver breakout used for UART-to-RS-485 conversion.* |

Table S4: Primary constituent ROV components. Column 1 includes links for more information. Prices retrieved September 29$^{th}$ 2026 and do not include sales tax, shipping, or tariff surcharges. “x” in the latter four columns represents inclusion of a component within an ROV version; “–” denotes optional equipment, i.e., navigational sensor suite items that can be utilized across $V1$–$V4$ depending on mission objectives. Manufacturer part numbers reflect the manufacturer or vendor part numbers listed on the linked product pages when available; “—” indicates that a clear part number was not listed or is configuration-specific. DeepSea unit cost includes cable assembly and bracket ($2,470 + $380 + $25); DeepSea and GNSS Compass prices are vendor quotes. The Kraken Solar Flare Mini 18000 is discontinued.

| **Item** | **Qty.** | **MFN part no.** | **Unit cost** | $V1$ | $V2$ | $V3$ | $V4$ |
|---|---|---|---|---|---|---|---|
| BlueROV2 | 1 | BR-100415 | $4,900 | x | x | x | x |
| BR Heavy upgrade kit | 1 | BR-101130 | $900 | x | x | x | x |
| GoPro 12 | 2 | CHDHX-121 | $369.99 | x | x | x | x |
| Dell Rugged laptop | 1 | configuration-specific | $4,000 | x | x | x | x |
| Pelican case | 1 | IM2950-00001 | $404.95 | x | x | x | x |
| BR tether spool | 1 | TMS-ASM-KIT-VP | $820 | x | x | | |
| BR Lumen Lights | 1 | BR-100737 | $770 | x | | | |
| BR Ping1D | 1 | BR-101684 | $430 | x | | | |
| Kraken Solar Flare 18k | 4 | SolarFlare-Mini18 | $850 | | x | x | |
| Payload Skid | 1 | BR-100233 | $320 | | x | x | x |
| Outland Topside Power Supply | 1 | OTPS1KW-R1-VP | $13,230 | | | x | x |
| Outland Tether Spool | 1 | RL-750 | $3,950 | | | x | x |
| DeepSea SeaStrobe | 4 | LSL-2075 | $2,875 | | | | x |
| WL DVL A50 | 1 | BR-101756 | $8,710 | – | – | x | x |
| WL UGPS | 1 | WL-11001-2 | $10,520 | – | – | x | x |
| GNSS Compass | 1 | GC2-POE / GC2-SER | $2,500 | – | – | x | x |

#### ROV camera settings and survey imagery

We initially utilized downward-facing GoPro HERO 10's to gather 4K video imagery using the settings listed below. However, due to image quality degradation upon extracting still images from these videos (see, e.g., Fig. S9), we discontinued the collection of downward-facing video, except to generally document the site. This switch was due to the downward-facing extracted stills exhibiting slight blurring, lack of definition, and lack of clear edges between, e.g., distinct categories of red or brown algae, or substrate. This is especially apparent when comparing the still images from Fig. S9 to the photographs in main text Fig. 5. While the extracted stills were generally sufficient for a human eye to make an identification, they proved to be limiting for the machine learning automation of extracting data regarding red, green, brown, and encrusting algae, and sessile and substrate type, from the still images — the details of which are beyond the scope of this field-methods focused manuscript.

For the primary downward-facing ROV survey camera, we switched to shooting 27.3 MP GPR (and JPEG preview) photographs via GoPro HERO 12, as the 12 (and newer) models allowed us to access GoPro Labs and the associated customized GoPro settings. Following collection, all JPEG and GPR image files were renamed to match the EXIF metadata timestamp recorded by the GoPro at capture time via `rename_jpg_gpr_EXIF.py`. Using `meter_mark.py`, images matching the survey time period were redistributed to transect folders for post-processing. Since 2026, `rov_imagery_processing` performs both steps as images are imported from the SD card, and can additionally select one image per user-defined distance traveled (Section S4).

Transect sets of downward-facing GPR photos (typically 50–200 photos) were batch edited in Adobe Lightroom Classic. This workflow included standardized edits as follows: a $4606 \times 4030$ crop to reduce corner distortion, utilizing the "remove chromatic aberration" tool, and applying denoise at 50%. Since 2026, `rov_imagery_processing` applies these standardized edits automatically and exports 16-bit TIFFs. Further adjustments to white balance, tone, and presence settings were applied individually to each image in the set. We retained shooting forward-facing 4K video for the forward-facing camera, as extracted stills were sufficient for the general identification of taxa such as demersal fish and conspicuous invertebrates (Fig. S10).

**4K video settings**:

- Mode = Video;
- Resolution = 4k;
- FPS = 30;
- Lens = Wide;
- Bit Rate = High;
- Shutter = Auto;
- White Balance = Native;
- ISO Min–Max = 100–3200;
- Color = Flat;
- Sharpness = Low.

**27.3 MP GPR photo settings**:

- Mode = Photo;
- Interval = 3 s;
- Lens = Wide;
- Output = RAW (GPR);
- EV Comp = 0;
- White Balance = Native;
- ISO Min–Max = 100–200;
- Sharpness = Low;
- Color = Flat;
- Shutter Speed = $\frac{1}{300}$ s.

To show image quality across whole transects rather than in selected frames, Table S5 links four complete transects of downward-facing imagery: one over flat seafloor and one along breakwater riprap for each of the $V1$ and $V2/V3$ configurations, with each pair flown at the same site.

Table S5: Complete transects of downward-facing survey imagery from the $V1$ and $V2/V3$ configurations (main text Table 1), for comparison with the selected frames in Fig. S9 and main text Fig. 5. All four transects are available at `https://osf.io/9dkp8/`, one folder per row, named by configuration, habitat, and date (e.g., `V1_flat_2022-08-15`). Because OSF can be slow to display individual images, we recommend downloading the imagery as zip files instead, either all four folders at once (direct download, 1.2 GB) or one folder at a time from the project's file list. Each pair was flown at the same site (the two flat-seafloor transects within 20 m of each other, the two riprap transects within 60 m); both sites are in Elliott Bay, Seattle, Washington. $V1$ imagery consists of stills extracted from 4K video, lit by 6,000 lm of Lumen lights; $V2/V3$ imagery consists of 27.3 MP raw (GPR) photographs lit by 57,600 lm of Kraken lights, developed with the standardized Lightroom edits described above, and subsampled to approximately one photograph per meter of transect. Transects were chosen as representative of each configuration and habitat; the $V2/V3$ riprap transect was chosen among those flown in `surftrak`. Depth and altitude are medians of 1 Hz telemetry over the transect (8.5–11 min each), with altitude from the Ping1D echosounder ($V1$; Table S4) or the DVL ($V2/V3$). The $V1$ transects, from our first surveys, predate the DVL and `surftrak` and were flown with the pilot controlling altitude; $V2/V3$ transects were flown entirely in `surftrak`, with the percentage of time within ±0.20 m of the target altitude in parentheses.

| **Configuration** | **Habitat, site, date** | **Depth (m)** | **Altitude (m), median (IQR)** | **Altitude control** | **Images** |
|---|---|---|---|---|---|
| $V1$<br>4K video stills | Flat seafloor<br>Centennial Park<br>2022-08-15 | 5.1 | 1.00 (0.89–1.09) | Pilot | 86 stills<br>3840 × 2160 |
| | Breakwater riprap<br>Elliott Bay Marina<br>2022-10-10 | 6.1 | 1.10 (0.94–1.26) | Pilot | 64 stills<br>3840 × 2160 |
| $V2/V3$<br>Raw photographs | Flat seafloor<br>Centennial Park<br>2024-10-08 | 4.2 | 0.68 (0.65–0.70) | `surftrak`<br>(100%) | 63 photos<br>4606 × 4030 |
| | Breakwater riprap<br>Elliott Bay Marina<br>2025-01-28 | 8.4 | 0.77 (0.71–0.84) | `surftrak`<br>(95%) | 60 photos<br>4606 × 4030 |

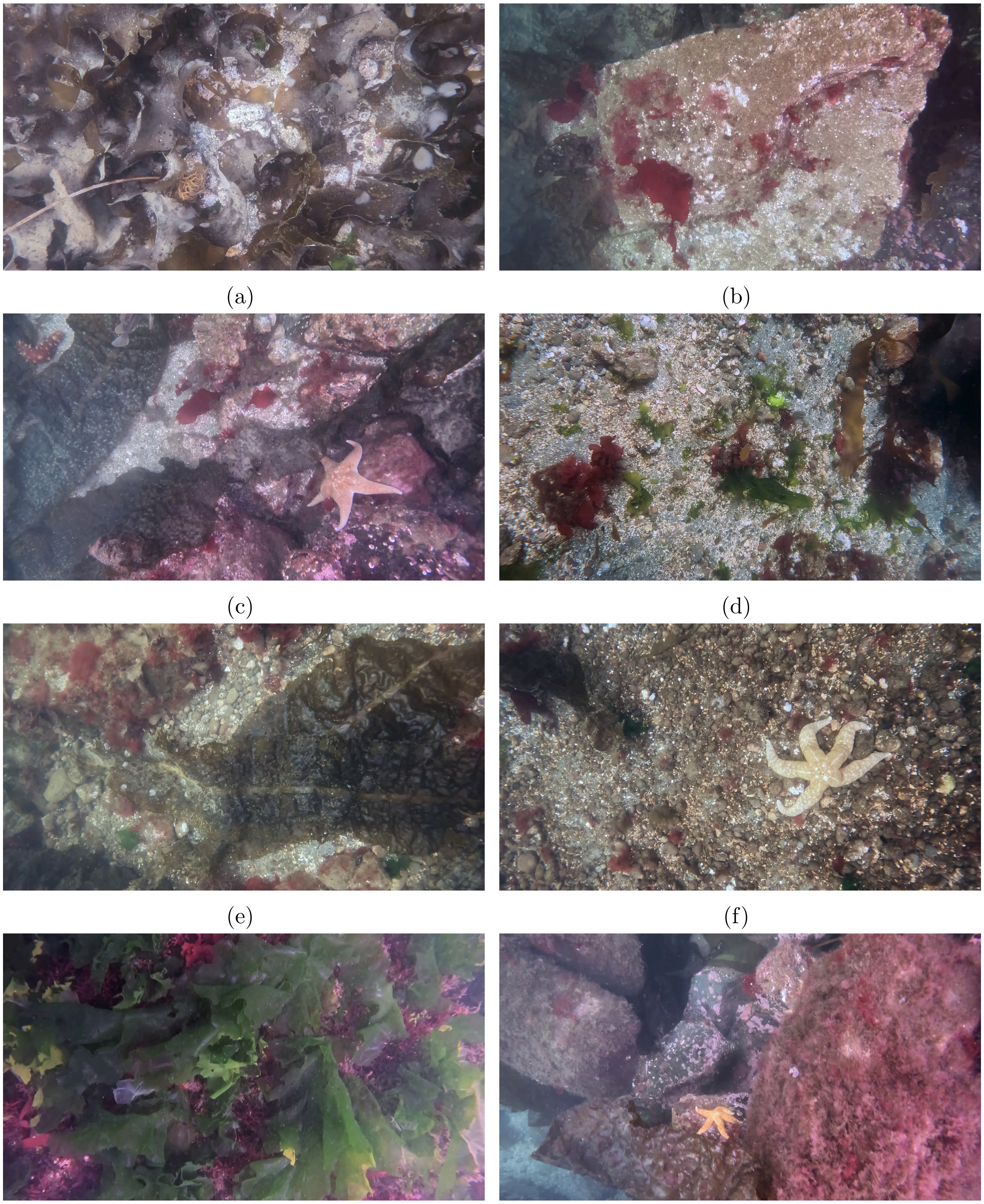

(a) (b) (c) (d) (e) (f) (g) (h)

Figure S9: Still images extracted from 4K, mp4 videos from our 2022 and 2023 ROV survey imagery. The degradation in quality is visible in the slight blurring, the lack of definition between categories, and the lack of clear edges between substrate types. Compare these images to the photographs in main text Fig. 5.

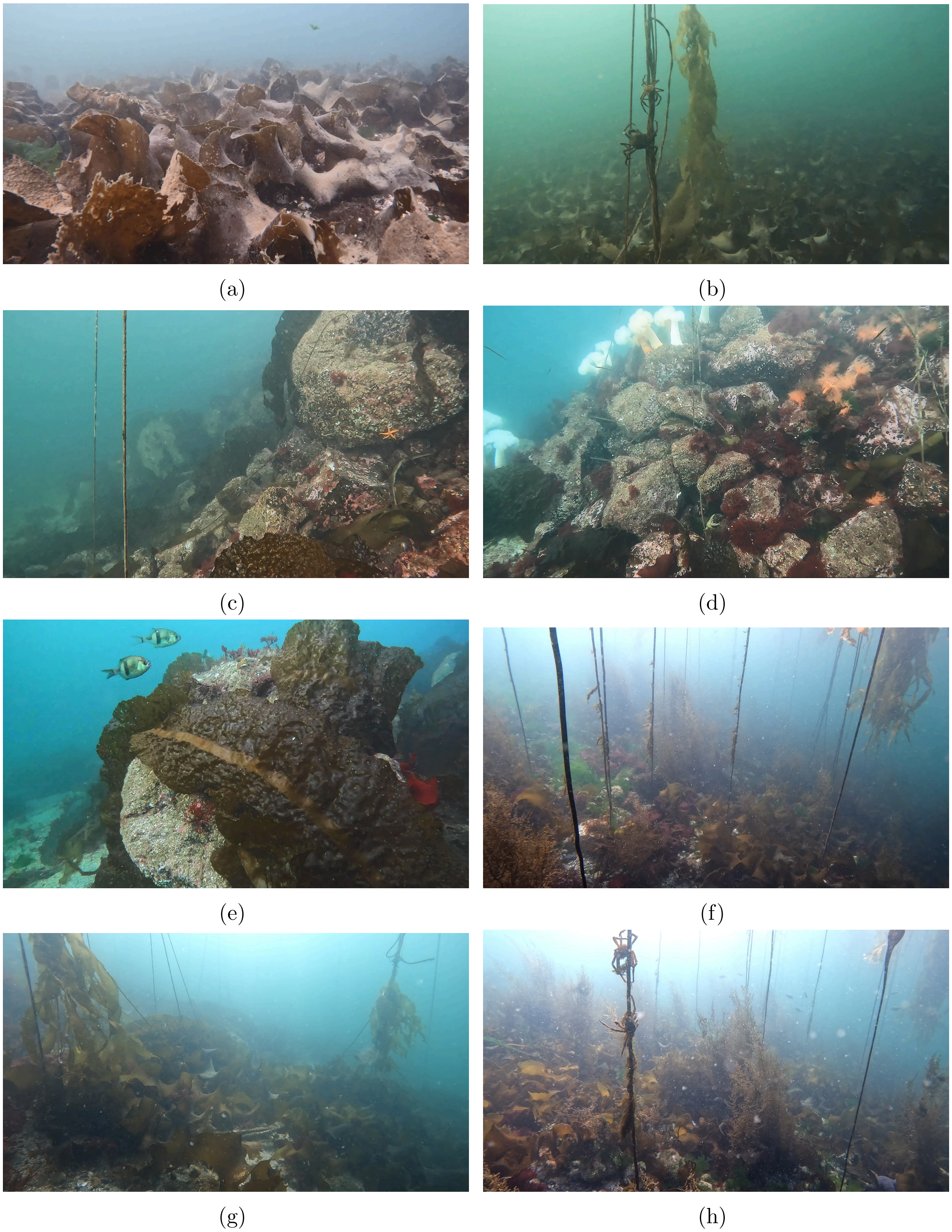

Figure S10: (a)-(h) Camera stills extracted from the forward-facing GoPro camera's 4K video. All imagery originate from Elliott Bay, Seattle, Washington. While there is still loss in quality with extracted stills, less fine-scale features and textures are required for basic species identification of mobile invertebrates, e.g., Kelp Crabs (*Pugettia producta*) and fish, e.g., Pile Perch (*Rhacochilus vacca*).

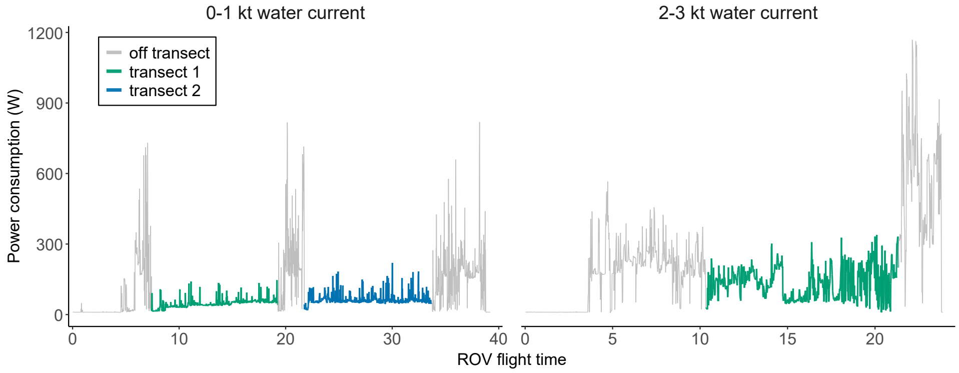

(a)

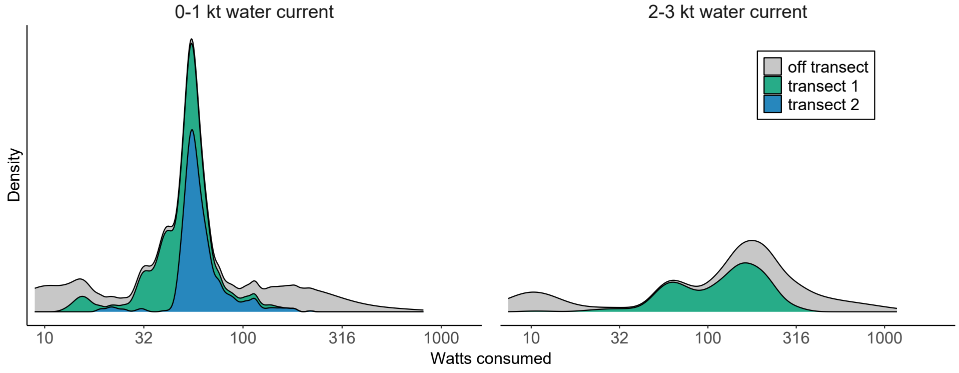

(b)

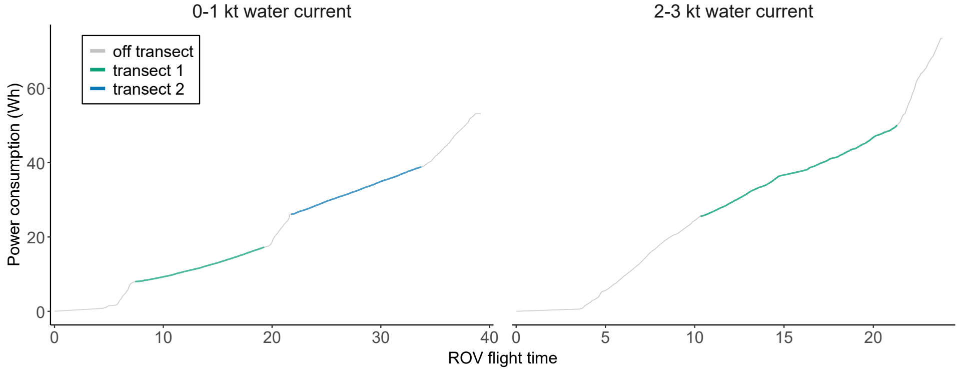

(c)

Figure S11: Power consumption values for the $V1$ configuration, with 0–1 kt of water current shown in the left column, and 2–3 kt of water current shown in the right column. The high-current flight was south of Waadah Island in the western Strait of Juan de Fuca, where water moved at 2–3 kt even near slack tide. Holding course required sustained thruster output of 50–70%, which heavily taxed the BR battery: power rose from approximately 60–100 W in low current to approximately 280 W, with cumulative energy use of 2.2 $\mathrm{Wh\,min^{-1}}$ on transect and 9.2 $\mathrm{Wh\,min^{-1}}$ during the more difficult surface recovery. Battery voltage dropped rapidly, total flight time was approximately 25 minutes — less than half a typical low-current flight — and the vehicle ultimately had to be pulled back by the tether, which the strain relief handled successfully. Flying against the current at the start of a mission, so that it assists recovery, can mitigate this where reef geometry and survey design allow.

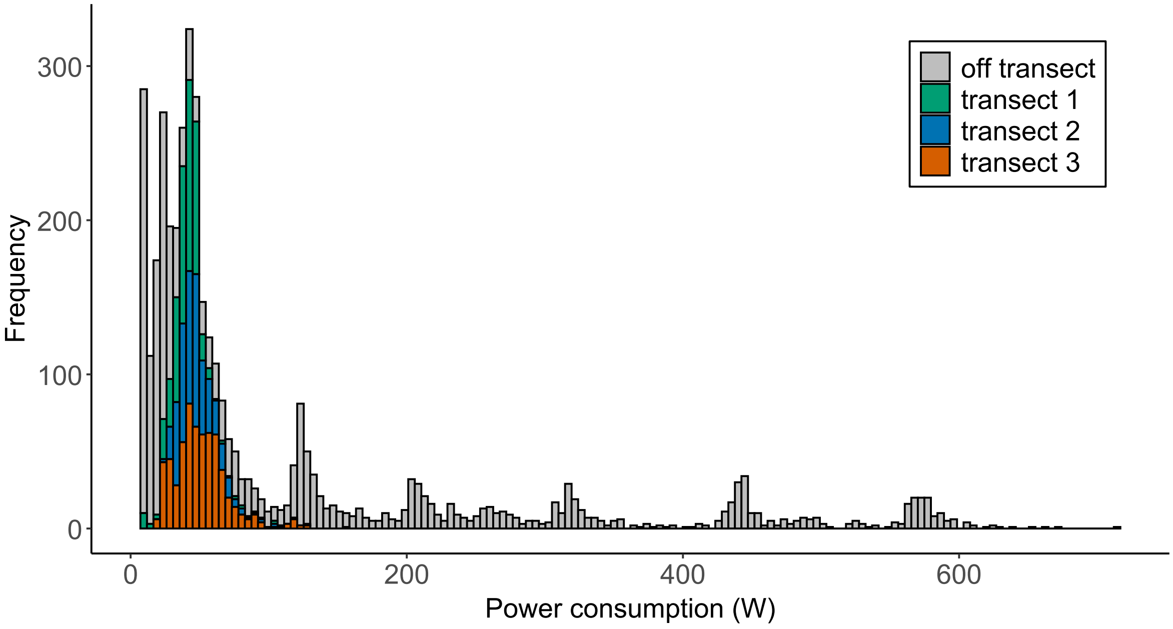

(a)

Figure S12: The natural scale i.e., non $\log_{10}(x+1)$ transformed data, exhibited in Fig. 4 for instantaneous power consumption in Watts (W).

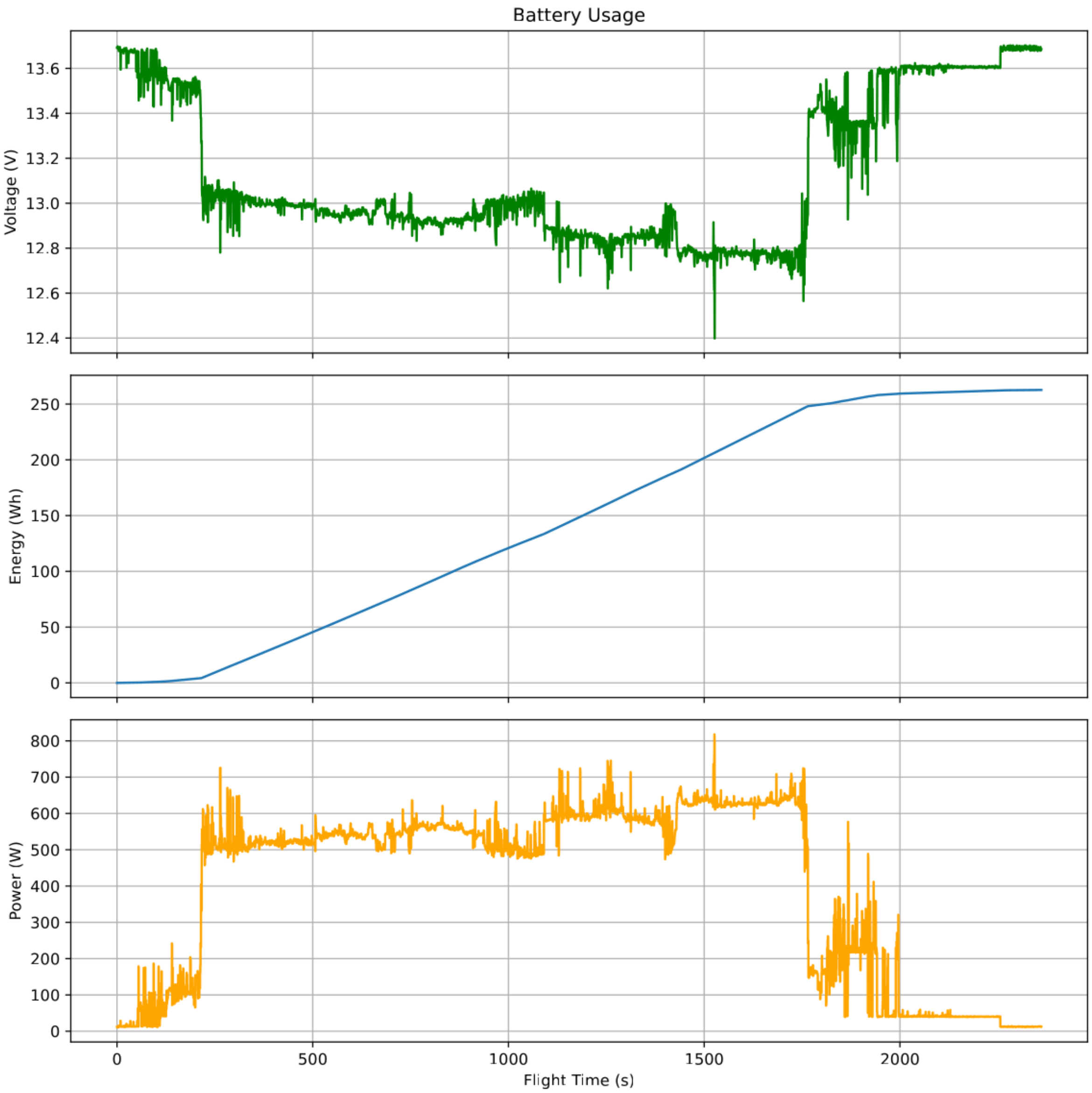

(a)

Figure S13: Power consumption from the $V4$ configuration with voltage (V), cumulative energy consumption (Wh), and power (W). The four DeepSea lights each consumed approximately 100 W of power, yielding 500 W baseline power consumption when the DeepSea lights are powered to 100% and the thrusters are around 20–30%. Increasing the thruster gain by increments of 10% increased power consumption by approximately 50 W, such that 50% thruster gain was found to associate with approximately 650 W power consumption. Despite running the thrusters consistently at "full throttle" for these tests, there are occasional spikes in power consumption, including a spike up to 800 W when at 50% thruster gain, close to the 1000 W limit of the OTPS system.
